# Distinct function of Chlamydomonas CTRA-CTR transporters in Cu assimilation and intracellular mobilization

**DOI:** 10.1101/2023.10.19.563170

**Authors:** Daniela Strenkert, Stefan Schmollinger, Srinand Paruthiyil, Bonnie C. Brown, Sydnee Green, Catherine M. Shafer, Patrice Salomé, Hosea Nelson, Crysten E. Blaby-Haas, Jeffrey L. Moseley, Sabeeha S. Merchant

## Abstract

Successful acclimation to copper (Cu) deficiency involves a fine balance between Cu import and export. In the unicellular green alga *Chlamydomonas reinhardtii*, Cu import is dependent on <u>C</u>opper <u>R</u>esponse <u>R</u>egulator <u>1</u> (CRR1), the master regulator of Cu homeostasis. Among CRR1 target genes are two Cu transporters belonging to the CTR/COPT gene family (*CTR1* and *CTR2*) and a related soluble cysteine-rich protein (CTR3). The ancestor of these green algal proteins was likely acquired from an ancient chytrid and contained conserved cysteine-rich domains (named the CTR-associated domains, CTRA) that are predicted to be involved in Cu acquisition. We show by reverse genetics that Chlamydomonas CTR1 and CTR2 are canonical Cu importers albeit with distinct affinities, while loss of CTR3 did not result in an observable phenotype under the conditions tested. Mutation of *CTR1*, but not *CTR2*, recapitulate the poor growth of *crr1* in Cu-deficient medium, consistent with a dominant role for CTR1 in high affinity Cu(I) uptake. Notably, the over-accumulation of Cu(I) in Zinc (Zn)-deficiency (20 times the quota) depends on CRR1 and both CTR1 and CTR2. CRR1-dependent activation of *CTR* gene expression needed for Cu over-accumulation can be bypassed by the provision of excess Cu in the growth medium. Over-accumulated Cu is sequestered into the acidocalcisome but can become remobilized by restoring Zn nutrition. This mobilization is also CRR1-dependent, and requires activation of *CTR2* expression, again distinguishing CTR2 from CTR1 and is consistent with the lower substrate affinity of CTR2.

## INTRODUCTION

Copper (Cu) is an essential element for all aerobic organisms where it participates in redox reactions or reactions involving oxygen chemistry [1]. This reactivity can be harmful to intracellular macromolecules. Furthermore, the affinity of Cu ions for metal binding sites in proteins can result in their mis-metalation [2]. Therefore, cells regulate Cu content very tightly. A balance between Cu assimilation and export is one common route to maintaining cellular Cu homeostasis. The *Enterococcus hirae* system, involving a *cop* operon encoding both influx and efflux P-type ATPase transporters, is a well-studied prototype for the maintenance of Cu homeostasis in prokaryotic systems [3]. In eukaryotes, Cu is taken up in two highly conserved steps: extracellular reduction of Cu(II) to Cu(I) followed by uptake via Cu(I) specific transporters [4–10].

The prototype assimilatory Cu(I) transporter, copper transport 1 (ctr1p), was originally discovered in yeast (*Saccharomyces cerevisiae*) with related proteins discovered in humans, plants, other fungi and algae by homology or by functional complementation of a yeast *ctr1* mutant. These transporters are called either CTR or COPT-type proteins (<u>C</u>opper <u>TR</u>ansporter in prokaryotes and mammals, <u>COPT</u> for <u>COP</u>per <u>T</u>ransporter in land plants). Typically, CTR/COPT family members have three trans-membrane helices (TM1-TM3), an N-terminal region that is rich in methionine (Met) residues, and a cysteine (Cys) rich, C-terminal region. Individual polypeptides assemble to form trimeric pores [11–14]. TM2 and TM3 contain the highly conserved motifs, MxxxM and GxxxG, respectively [15, 16]. The M-rich motifs are extracellular and essential for Cu(I) transport activity: they bring Cu(I) to the entrance of the homotrimeric pore. The pore itself is composed of methionine triads and serves to channel Cu across the lipid bilayer. After Cu sequestration through the pore, Cu(I) binds to the C-terminal Cys-rich motifs of CTR, where it is delivered to soluble metallo-chaperones for metalation of cuproproteins and intracellular distribution. The association of intracellular Cu(I) with Cu chaperones enables passive transport of Cu(I).

Where tested, the CTRs localize to the plasma membrane of eukaryotic cells and mediate high affinity Cu(I) uptake: their loss of function can lead to cell death since Cu(I) is an essential element for most aerobic life [1]. Genes encoding Cu transporters are typically transcriptionally activated in response to poor Cu nutrition, enabling Cu acquisition in such a situation [17, 18].

The Arabidopsis (*Arabidopsis thaliana*) genome encodes six members belonging to the CTR-type family of Cu transporters (COPT1-COPT6) [19]. Of these, COPT1, COPT2 and COPT6 are plasma membrane-localized, fully rescuing the yeast *ctr1* mutant, and are involved in Cu uptake into the cell [20–23]. COPT5, on the other hand, exports Cu(I) from intracellular storage sites (or vacuoles) [24, 25]. Roles for COPT3 and COPT4 have yet to be defined.

The *Chlamydomonas reinhardtii* genome encodes four CTR/COPT members: CrCTR1 and CrC-TR2, which the classic 3 trans-membrane topology, CrCTR3, which is derived from a gene duplication of *CTR2* but has lost the hydrophobic trans-membrane domains, and CrCOPT1, which shows the 3 trans-membrane structure but lacks the amino-terminal region that is rich in Met and Cys residues [7]. The distinct nomenclature reflects the closer relationship of CrCTR1 and CrC-TR2 to fungal proteins and COPT1 to plant proteins. The pattern of expression of the four corresponding genes in response to Cu nutrition suggests a role for the CTRs, but not for CrCOPT1, in high affinity Cu(I) assimilation [7]. Specifically, *CTR* transcripts are increased in abundance under poor Cu nutrition but reduced in the Cu excess situation. The regulation occurs at the level of transcription through associated

Cu response elements (CuREs) and a transcription factor, <u>C</u>opper <u>R</u>esponse <u>R</u>egulator <u>1</u> (CRR1). CRR1 binds to CuREs through a green-lineage specific DNA binding domain. Loss of function of CRR1 results in the loss of expression of ∼ 63 genes (that define the nutritional Cu regulon in Chlamydomonas), including the *CTR*s, and hence causes poor growth in Cu-deficient medium. *C. reinhardtii* CTR-like proteins are in other green algae but are more similar to fungal proteins as compared to the proteins found in land plants. Unique characteristics of these proteins are the replacement of some of the Met-residues in the N-terminal Cu(I)-binding region by Cys-residues. Chlamydomonas *CTR1* and *CTR2* rescue the yeast *ctr1* strain underscoring their function in Cu(I) transport [7]. CTR3 is a soluble protein [26]: it may play a role in high affinity Cu uptake by recruiting Cu(I) to the assimilation pathway components, analogous to the proposed function of the soluble FE ASSIMILATION (FEA) proteins during iron (Fe) assimilation in *C. reinhardtii* [27]. In previous work, we investigated Cu metabolism and homeostasis in *C. reinhardtii* in response to Cu nutrition [28]. Although the genome potentially encodes dozens of cuproproteins, three proteins are dominant in contributing to the Cu quota of the algal cell: plastocyanin in the chloroplast, cytochrome oxidase in the mitochondrion, and a ferroxidase (FOX1) involved in high affinity Fe uptake at the plasma membrane. In Fe-replete medium, *FOX1* expression can be repressed, leaving only two proteins as biomarkers of intracellular Cu status.

Plastocyanin is the most abundant cuproprotein in *C. reinhardtii* (which is functionally equivalent to a photosynthetic cell in plants), but in many algae its function in photosynthesis can be replaced by a heme protein, cytochrome *c*_6_ (Cyt *c*_6_). The expression of the corresponding *CYC6* gene is activated by CRR1 under Cu-deficiency [29, 30]. Accordingly, plastocyanin is on the bottom of the hierarchy for Cu distribution in *C. reinhardtii*, and is a sensitive indicator of intracellular Cu status. Importantly, the function of cytochrome oxidase in respiration cannot be covered by any other protein and it therefore receives Cu with the highest priority: cytochrome oxidase abundance is decreased only in the most severely Cu-deficient situation. On the other end of the spectrum, when Cu(I) uptake is in excess and cells over-accumulate Cu(I) species, the metal is stored in the acidocalcisome, most likely in association with Cys or related metabolites [31].

In this work, we used a reverse genetic approach to test and distinguish the functions of the *C. reinhardtii* CTR proteins in the context of cuproprotein assembly and Cu homeostasis. By measuring cellular Cu content chemically, monitoring individual Cu protein abundances to assess intracellular Cu status, and visualizing a key Cu(I) storage site, we conclude that CTR1 and CTR2 are both required for Cu assimilation although with different characteristics, while CTR2 is exclusively required for Cu efflux from intracellular storage sites. CTR3 is an extracellular periplasmic protein that appears to be dispensable under laboratory conditions.

## RESULTS AND DISCUSSION

### Cu uptake is impaired in the *crr1* mutant

Since the *CTR* genes are targets of the CRR1 regulon (Figure 1A), we tested a *crr1* mutant and its complemented *CRR1* strain for Cu uptake. To this end, we first grew both strains in Cu-free medium to deplete intracellular Cu stores and added 2 µM Cu(II), provided as CuEDTA, to each culture. We collected cells 1 h before Cu(II) addition, and 0.5 and 3 h after Cu(II) addition (see Figure 1B) to measure cellular Cu content by Inductively coupled plasma-tandem mass spectrometry (ICP-MS/MS). We observed that both strains have very low Cu levels following growth under Cu-deficient conditions (Figure 1C). Importantly, while Cu content increased in the complemented *CRR1* strain rapidly (0.5 h time point), the *crr1-2* mutant strain accumulated very little Cu. Indeed, the *crr1* mutant reached less than 10% of the Cu amount in the complemented *CRR1* line after 3 h in the presence of external Cu supply (Figure 1C), consistent with a role for CRR1-dependent inducible Cu uptake in Chlamydomonas, possibly via one or more of the CTR transporters [8].

**Figure 1:**
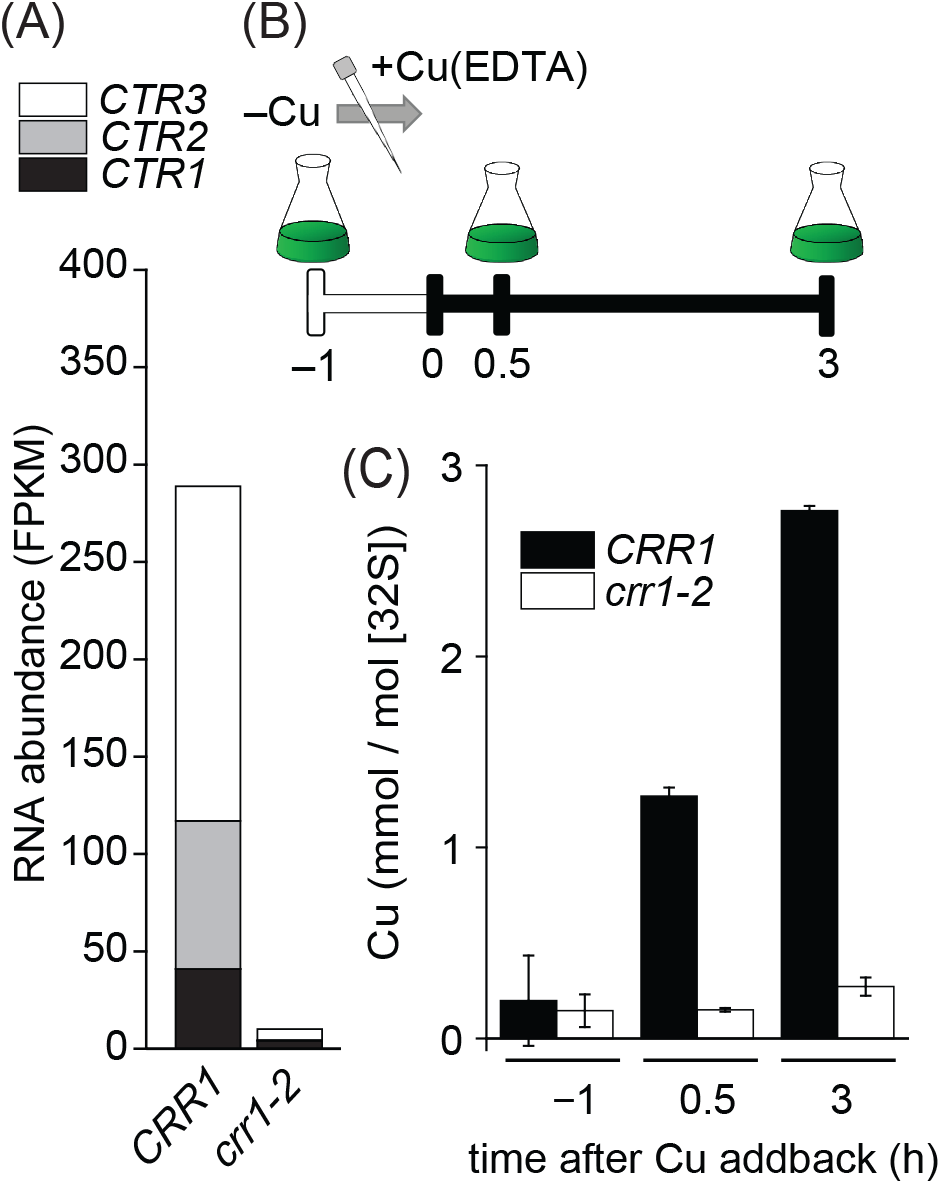
CRR1 is required for *CTR* gene expression and for Cu uptake. (A) *CTR1*, *CTR2* and *CTR3* mRNA abundances (in FPKMs) in Cu deficient c*rr1-2* (mutant) and *CRR1* (wildtype) grown as indicated, according to (Castruita et al. 2011). (B) Experimental design for Cu resupply experiments. Cells were grown in medium without Cu supplementation until early exponential growth. 2 µM Cu(EDTA) was added at time 0. (C) ICP-MS/MS analysis at the indicated time points. Cu content, normalized to ^32^S as a measure for biomass, before (−1h) and after (0.5h and 3h) Cu addition. Shown are data points, averages and StDEV of three independent experiments.

### Evolution of CTR proteins from Chlamydomonas

Both CTR1 and CTR2 of Chlamydomonas have an extracellular N-terminal region enriched with Cys residues, an intracellular C-terminal region with Cys and/or His motifs, and three pore-forming TM domains for Cu(I) transport characteristic of the CTR family (Supplemental Figures 1 and 2). In addition to the Cys residues that are expected to bind Cu on the extracellular side of the membrane, Mets motifs (MxxM or MxM) are also present; 5 Mets motifs in CTR1 and 6 in CTR2 (Supplemental Figures 1 and 2). In contrast, COPT1 has a relatively short N-terminal soluble region that lacks Mets motifs or Cys residues but is dominated by histidine residues (Supplemental Figure 3). Based on blastp alignments between CTR3 and the N-terminal regions of CTR1 and CTR2, we identified 2 separate Cys-rich regions in CTR1 and CTR2, with 3 such regions in CTR3 (Supplemental Figures 1,2 and 4). Based on percent identity, CTR3 is likely a duplication of the N-terminal region of CTR2 followed by a tandem duplication of region 2 within CTR3. Each identified region is predicted to fold into separate structural domains, with proline-rich regions separating the two domains in CTR2 and the three domains in CTR3 (Supplemental Figures 2 and 4). Based on this analysis, we have named this conserved domain the CTRA (CTR-associated) domain and refer to this subfamily of CTRs as the CTRA-CTR proteins.

A sequence similarity network (SSN) of CTR proteins (based on presence of IPR007274) and CTR3-like proteins (based on a jackhmmer search) reveals multiple major proteins clusters that are separated largely in accordance with taxonomy (Figure 2A). Chlamydomonas COPT1 and other closely related green algal CTR proteins are connected to the land plant CTRs, which are further divided into two subclusters, represented by AtCOPT5 and AtCOPT1 (Figure 2A). Two exceptions to the congruence between taxonomy and protein clusters are present. One cluster contains the CTR1/CTR2/CTR3 proteins from Chlamydomonas together with other green algal proteins, fungal proteins from Fungi incertae sedis, and proteins from Amoebozoa. The second cluster contains proteins from an assortment of lineages, including green algae and various protists. The protist cluster contains proteins from several Chlamydomonas species, but not *C. reinhardtii*. The CTR1/CTR2/CTR3 cluster contains a mix of stand-alone CTR3-like proteins and CTR proteins that contain one or more CTRA domains (Figure 2A).

**Figure 2:**
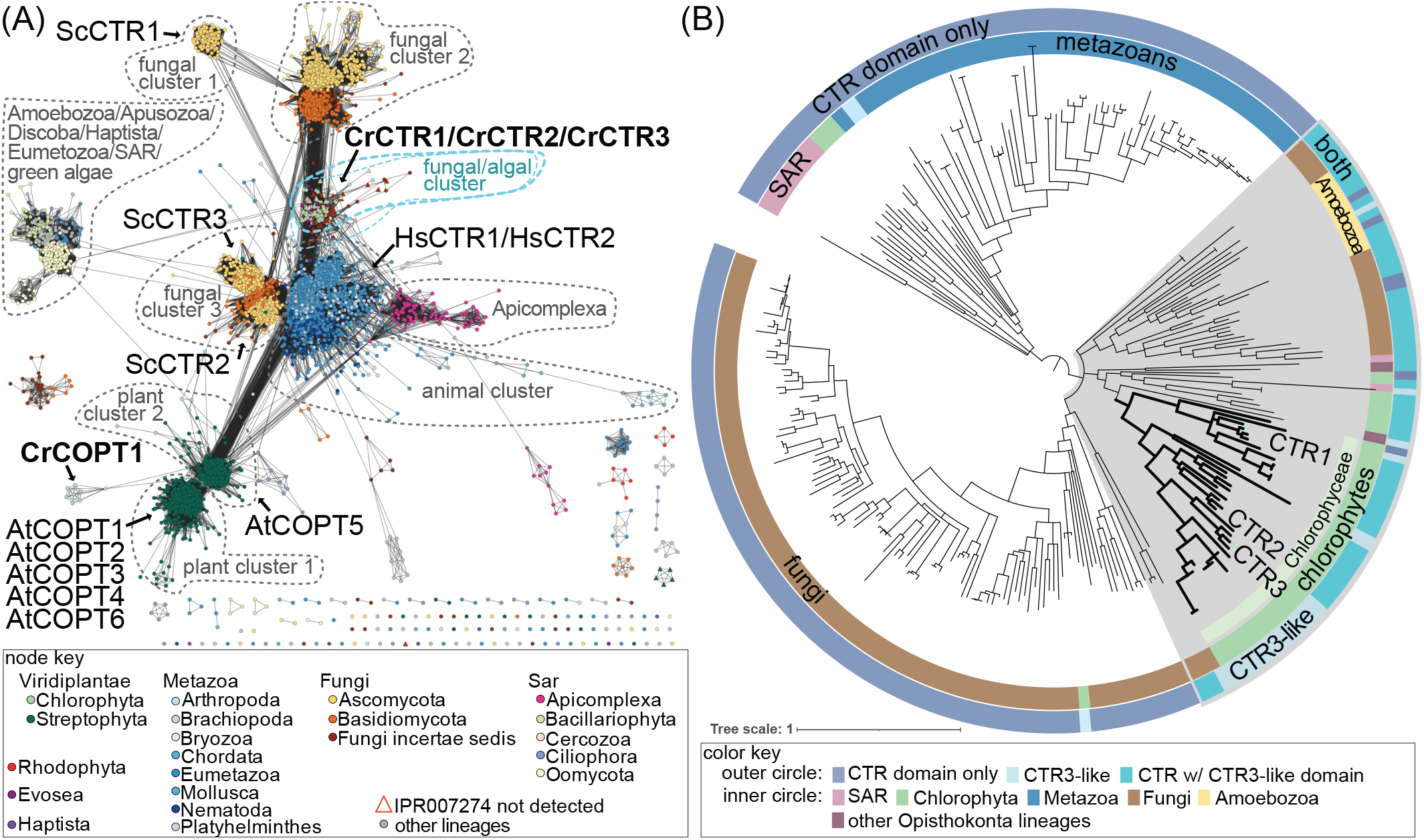
Clamydomonas CTR1, CTR2, and CTR3 belong to the CTRA-containing family. (A) Sequence similarity network on the CTR family and CTRA-containing proteins. Nodes are colored based on taxonomy according to the color key. The clusters to which the *Saccharomyces cerevisiae* (labeled with “Sc”), *Arabidopsis thaliana* (labeled with “At”), *Homo sapiens* (labeled with “Hs”), and the *Chlamydomonas reinhardtii* (labeled with “Cr”) CTR-family proteins belong are indicated with an arrow. The major distinct clusters and subclusters are delineated with a dotted line and labelled. The CTRA-containing cluster is outlined with a blue dotted line, and a close-up view is provided in the blue square in the upper right of the panel. (B) Phylogenetic tree of proteins similar to CTR1, CTR2, and CTR3. The outer circle is used to indicate whether the leaf represents a protein with a CTR domain (without the CTRA domain), a CTRA-CTR fusion, or a CTRA-containing protein (without the CTR domain). The inner circle is used to convey taxonomic information for each leaf according the to the color key. The Chlorophyceae-specific clade containing *C. reinhardtii* CTR1, CTR2, and CTR3 is highlighted with thicker branches and the location of these proteins is labeled.

This analysis also reveals that fusion proteins involving CTR are not unique to CTR1/CTR2. Five other types of N-terminal extensions involving annotated domains are present, as well as five types of C-terminal extensions (Supplemental Figure 5). Metal-related examples are the C-terminal HMA (Heavy Metal Associated) domains found in the protist cluster and the C-terminal BIM1 domains found in fungi. Small soluble proteins containing the HMA domain often function as Cu chaperones and are proposed to acquire Cu directly from CTR [32], while BIM1 is involved in Cu acquisition [33]. Since this analysis relies on identification of annotated domains, undescribed domains in addition to the CTRA domains found in CTR1/CTR2/CTR3 likely exist. One example is the unannotated N-terminal domain found in the protist cluster, represented by CTR protein from the oomycete *Albugo candida* (Supplemental Figure 5), which is also found as a stand-alone protein in other protists and green algae but is not related with the CTRA domain. Although this type of CTR fusion protein is not found in *Chlamydomonas reinhardtii*, the stand-alone protein is conserved (Cre05.g236039) and is a target of CRR1, suggesting that this uncharacterized protein may function in Cu acquisition.

Phylogenetic reconstruction of CTR1, CTR2, CTR3 and similar proteins confirms the relatedness of these proteins to CTRA-CTR proteins from Fungi incertae sedis and Amoebozoa. The phylogenetic tree suggests that a geen algal ancestor that existed before the split of Pedinophyceae, Chlorophyceae, and Trebouxiophyceae could have acquired a CTRA-CTR fusion protein from an ancestor of the Chytridiomycetes. Chytridiomycetes are typically known as aquatic parasites and can infect green algae, but some species form a facultative mutualistic relationship with green algae [34]. The evolution of CTR3, likely resulting from a partial duplication of the ancestral *CTR2*, appears to have occurred in an ancestor of the Chlamydomonadales. There is also evidence of independent duplications leading to the evolution of CTRA-containing proteins that lack the CTR TM helixes (Figure 2B).

Previous biochemical analyses indicated plasma membrane localization for CTR1 and CTR2, suggesting that one or both proteins might function in vivo in Cu(I) transport [7].

By contrast, CTR3 was previously detected in soluble fractions of cells rather than in the membrane, consistent with the absence of TM domains in this protein [26]. We asked whether CTR3 might be a secreted protein, in a manner analogous to the periplasmic FEA proteins suspected to be involved in Fe assimilation [27], by monitoring CTR3 abundance in the spent medium of a cell-wall deficient (*cw*) strain (Supplemental Figure 6). A protein that is located to the periplasmic space is lost to the medium in a *cw* strain. We concentrated the proteins in the spent medium by trichloroacetic acid (TCA) precipitation, followed by SDS-PAGE separation and immunoblotting with an antibody raised against CTR3. We detected the presence of CTR3 only in the spent medium of *cw* cultures under Cu-deficient conditions (Supplemental Figure 6). We validated the Cu status of the cells by immunoblotting for Cox2b (lower abundance in low Cu). We also detected FEAs in the spent medium. We conclude that CTR3 is likely not a transporter, nor does it have strong, stable association with a plasma membrane–localized protein (e.g. CTR1 or CTR2) to be retained by *cw* cells.

### Reverse genetics to assess the function of each member of the Chlamydomonas CTR family

To directly assess the function(s) of individual CTR proteins in Chlamydomonas, we generated or identified candidate loss of function mutants. For *CTR1*, we used CRISPR/Cpf1-based gene editing [35–37] (see Methods), while for *CTR2* and *CTR3*, we obtained candidate loss of function insertion mutants from the Chlamy-domonas Library Project, CLiP [38]. We introduced two in frame stop codons within the first exon of the *CTR1* gene (Figure 3A, grey filled arrow). From 178 transformants screened, we identified five independent lines, named *ctr1-1* through *ctr1-5*. Sequencing of the PCR-amplified product spanning the gene editing site confirmed the presence of the two stop codons in all five *ctr1* lines, with *ctr1-2* harboring an additional insertion of three in frame codons (Supplemental Figure 7). Molecular analysis of *ctr1* lines indicated the absence of *CTR1* transcripts (Figure 3B) and the absence of the corresponding polypeptide (Figure 3C). For *CTR2* and *CTR3*, we confirmed disruption of the loci by PCR and Sanger sequencing. While *ctr2-1* has an insertion in the second exon of *CTR2* and a complete loss of *CTR2* mRNA, we noted that *ctr2-2* has an insertion in the second intron and expresses a low, but detectable, amount of the *CTR2* transcript (Figure 3B, 3.2%). Immuno-detection confirmed that *ctr2-2* is a weak allele with a small amount of residual protein (Figure 3D). For *CTR3*, genotyping confirmed the location of the insert in *ctr3-1*, and qRT-PCR and immunodetection, respectively confirmed the absence of mRNA and protein (Figure 3B & C).

**Figure 3:**
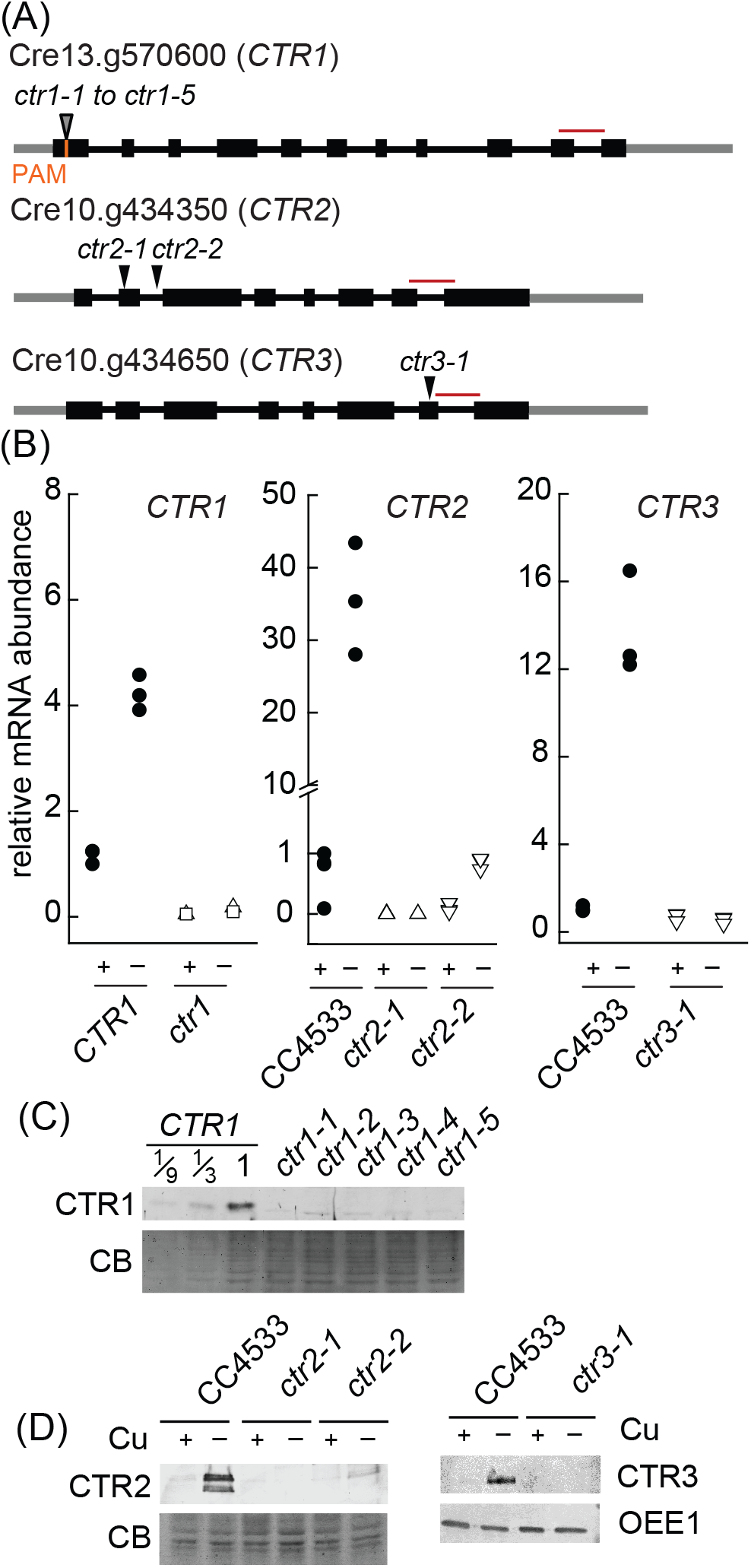
Molecular analysis of *ctr* mutants. (A) Location of inserts or target regions in all CTR gene models. The 3’ and 5’ untranslated regions are represented by a thin gray line, exons (black boxes) are numbered and thin black lines connecting exons are indicative of introns. The location of the amiRNA target site for *CTR1* (open arrowhead), the CPF1 target sequence (PAM, orange) and CPF1 mediated insertion site of two stop codons (grey arrowhead) and insertion sites for *CTR2* and *CTR3* (filled arrowheads) mutants are shown. Amplicons used for qRT-PCR in (B) are shown in dark red. (B) Relative transcript abundances of *CTR1*, *CTR2* and *CTR3* were determined using quantitative RT-PCR. Cells were grown in either Cu deficient (–) or replete (+) conditions as indicated. CRISPR reference lines (*CTR1*) are represented by filled circles, while open triangle up symbols indicate *ctr1-1* lines and open square symbols indicate *ctr1-3* lines, *ctr2-1* (open, up triangles) and *ctr2-2* (open down triangles) and *ctr3-1* (open down triangle). Each symbol represents an independent experiment. (C,D) 20 µg of total cell lysates were separated by SDS-PAGE (10% monomer), followed by immunoblotting using antisera against CTR1, CTR2 or CTR3, respectively. Coomassie blue (CB) or OEE1 serves as a loading control. Samples shown in (D) were from cultures grown in Cu deficient growth medium.

### CTR1 and CTR2 mediate Cu uptake in Cu replete cells

We measured the Cu content of each mutant and their wild-type strains (CC4533 for insertion mutants; CC425 for *ctr1* mutant strains) in medium containing sufficient (2 µM) but not excess Cu [28]. For *ctr1*, the Cu content reached 77% of wild-type levels (Figure 4A, shown is the average of all five allelic strains). Likewise, both the null *ctr2-1* strain and the weak *ctr2-1* strain had less Cu compared to the wild type, with *ctr2-1* showing a stronger decrease with respect to Cu content (66% of wild-type levels) than *ctr2-2* (77% of wild-type levels). We conclude that CTR1 and CTR2 both contribute to Cu assimilation in Cu-replete conditions. When we monitored the abundance of *CTR1* and *CTR2* transcripts in the mutants, we observed a dramatic 46-fold increase in *CTR2* transcripts in the *ctr1* mutants, and a smaller ∼ 3-fold increase for *CTR1* transcripts in the *ctr2-1* mutant and up to 2-fold increase in *ctr2-2* (Figure 4B). Thus, the absence of either transporter triggers induction of the Cu assimilation machinery, albeit to different degrees. In both cases, the induction of one transporter gene does not fully compensate for the loss of the other, speaking to distinct functions for CTR1 and CTR2.

**Figure 4:**
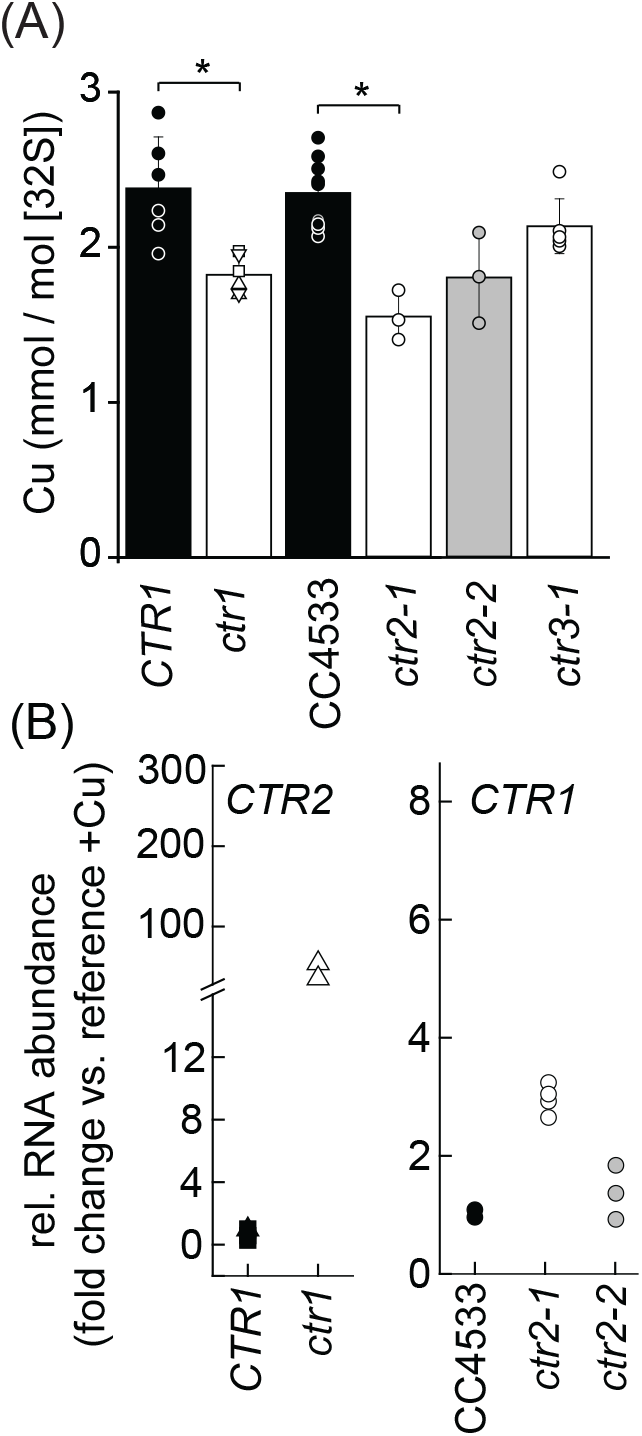
CTR dependent Cu uptake in Cu replete cultures. (A). Cu content from strains as indicated was determined by ICP-MS/MS. Black symbols and bars indicate the corresponding wild-type strains. Null mutants are indicated in white and a leaky mutant in grey as follows: *ctr1-1* (triangle up), *ctr1-2* (triangle down), *ctr1-3* (square), *ctr1-4* (circle), *ctr1-5* (diamond). Shown are data points, averages and StDEV of 3-9 independent experiments. (B) Relative transcript abundances of *CTR1* and *CTR2* were determined using quantitative RT-PCR. Cells were grown in Cu replete conditions. Samples collected from wild-type background strain (black triangle up), *ctr1* mutant lines (white triangle up). reference strain CC4533 (filled circles), *ctr2-2* (open, gray circles) and *ctr2-1* (open white circles). Each symbol represents an independent experiment.

We hypothesized that the observed increase in *CTR1* transcripts in *ctr2* cells under classic Cu-replete conditions (and *CTR2* transcripts in Cu-replete *ctr1* cells) relative to wild-type cells may reflect an intracellular Cu deficiency, which would lead to the activation of the CRR1 regulon. We tested this hypothesis by immunoblot analysis with the Cu deficiency marker Cyt *c*_6_. Indeed, we detected Cyt *c*_6_ in both *ctr2* mutants grown in TAP medium with 2 µM Cu (Cu-replete condition) (Supplemental Figure 8AB) with the level of Cyt *c*_6_ accumulation proportional to the magnitude of the Cu deficiency. By contrast, the *ctr1-1* mutant strain showed no evidence of internal Cu deficiency, as evidenced by the absence of Cyt *c*_6_ under Cu-replete conditions (Supplemental Figure 8A), suggesting that another mechanism is responsible for the compensatory increase of *CTR2* transcripts in the *ctr1* strain. How the changes in transcript levels translate to the abundance of the corresponding transporters, if any, has not been determined.

### Loss of CTR1 or CTR2 results in stronger phenotypes in Cu deficiency

Despite the lower Cu content of *ctr1* and *ctr2* mutants, we noticed no effect on their growth relative to their wild-type strains in Cu-replete conditions (Figure 5A, Supplemental Figure 8C). We therefore tested the mutants for growth in Cu-deficient medium when both genes are normally induced (Figure 5A, Supplemental Figure 8C). The *ctr2* and *ctr3* strains had doubling times comparable to those of the corresponding wild types, about 8 h in Cu-replete medium, with doubling times for *ctr2* mutant strains appearing to lag behind that of the wild type, although this difference did not reach significance (Supplemental Figure 8C). In sharp contrast, the *ctr1* strains were unable to grow in Cu-deficient medium recapitulating the *crr1* phenotype (Figure 5B). When we tested the *ctr1* strains for the accumulation of sentinel proteins for intracellular Cu status, plastocyanin and Cyt *c*_6_, we noted a normal pattern of Cu nutrition-dependent accumulation for plastocyanin and Cyt *c*_6_, indicating that the CRR1 regulon functions properly in the *ctr1-1* mutant (Supplemental Figure 8A). This observation suggests that the very poor growth displayed by *crr1* may be attributed to the lack of induction of *CTR1* in Cu-deficient conditions (Figure 5B). The *ctr2* strains also showed wild-type accumulation of plastocyanin and Cyt *c*_6_ in Cu-deficient medium, confirming, together with the absence of a growth phenotype, successful acclimation to Cu deficiency (Supplemental Figure 8B). The *ctr2-1* strain, which is more Cu-deficient, accumulated more Cyt *c*_6_ than did the *ctr2-2* strain in Cu replete conditions.

**Figure 5:**
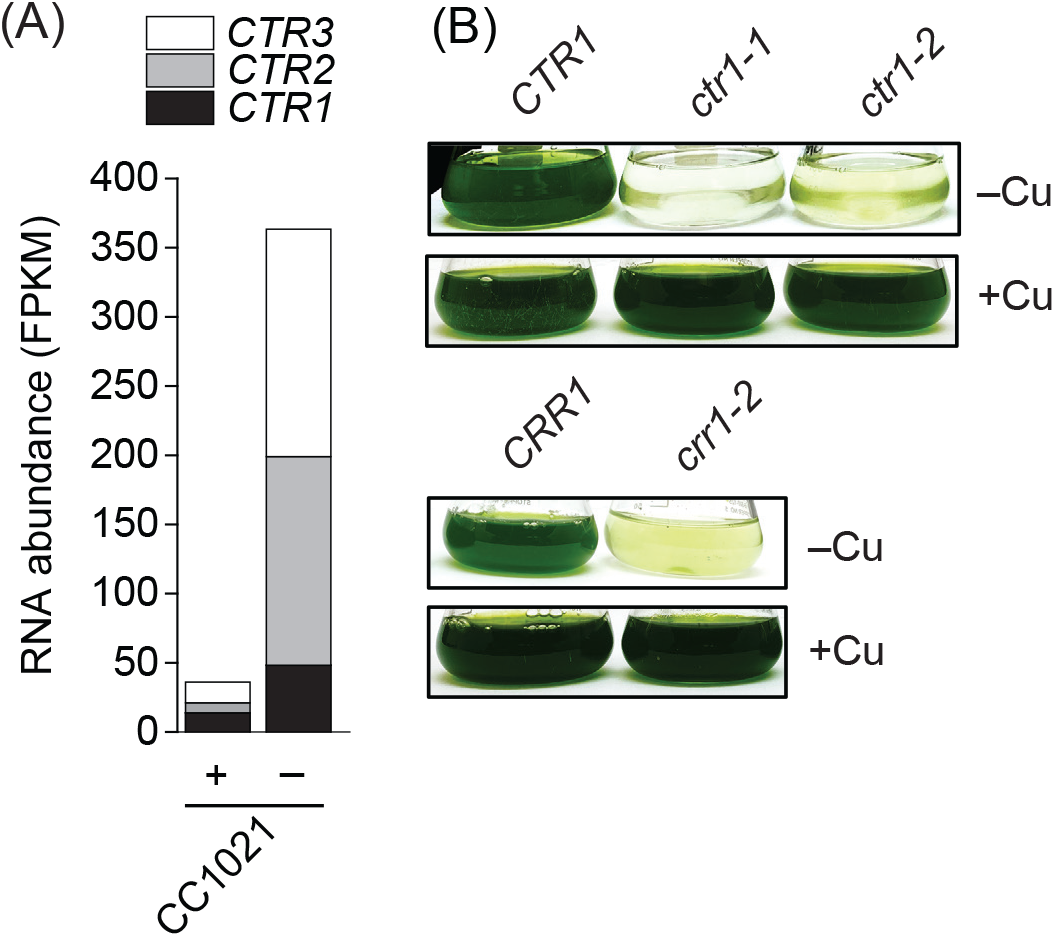
Although both CTR1 and CTR2 are up-regulated under poor Cu nutrition, only *ctr1* mutants recapitulate the Cu nutrition-dependent *crr1* phenotype. (A) *CTR1*, *CTR2* and *CTR3* mRNA abundances (in FPKM) in the Cu deficient wildtype strain CC1021 (Castruita et al. 2011). (B) Strains were grown photoheterotrophically under Cu-deficient (–) or Cu-replete (+) conditions. Pictures of flasks were taken six days post-inoculation.

We conclude that the phenotypes of the *ctr1* and *ctr2* strains under poor Cu nutrition are distinct and reinforces the notion that they have non-redundant roles in Cu homeostasis.

### Distinct functions of CTR1 and CTR2 in high- and low-affinity Cu uptake

To directly assess the function of CTRs in Cu(I) uptake, we measured the total cellular Cu content of Cu-deficient cells by ICP-MS/MS either before (1 h, to establish the ground state) or after (+0.5 h and 3 h) addition of 100 nM Cu to probe high-affinity uptake. We also tested low-affinity Cu uptake by performing the same experiment with the addition of 2 µM Cu (Figure 6). With the addition of 2 µM Cu, the Cu uptake of the *ctr1* and *ctr3* mutants was indistinguishable from that of their corresponding wild-type reference strains (Figure 6A and Supplemental Figure 9). Notably, the two *ctr2* mutant strains showed poor uptake kinetics (Figure 6D). Even at 3 h after addition of 2 µM Cu, the Cu content of the *ctr2-1* and *ctr2-2* strains was only 14% and 20% of the reference wild type, respectively, indicating that CTR2 is responsible for a large portion of the Cu assimilated at higher extracellular Cu concentrations. This result is consistent with the apparently normal Cu uptake seen in the *ctr1* mutant strains at high extracellular Cu, with Cu uptake occurring through CTR2. When we assessed high-affinity transport with the addition of 100 nM Cu, the *ctr1* mutants had 10-fold less Cu than the corresponding wild type (Figure 6B), explaining its poor growth in Cu-deficient conditions (Figure 5B). The *ctr2* mutants accumulated Cu, albeit less than the wild type (Figure 6E). We speculate that the slower but continuous accumulation of Cu in the *ctr2* strains growing in the presence of 100 nM Cu, presumably through CTR1, allows growth of *ctr2* in Cu-deficient medium, and indicates that CTR1 contributes predominantly to high-affinity Cu uptake.

**Figure 6:**
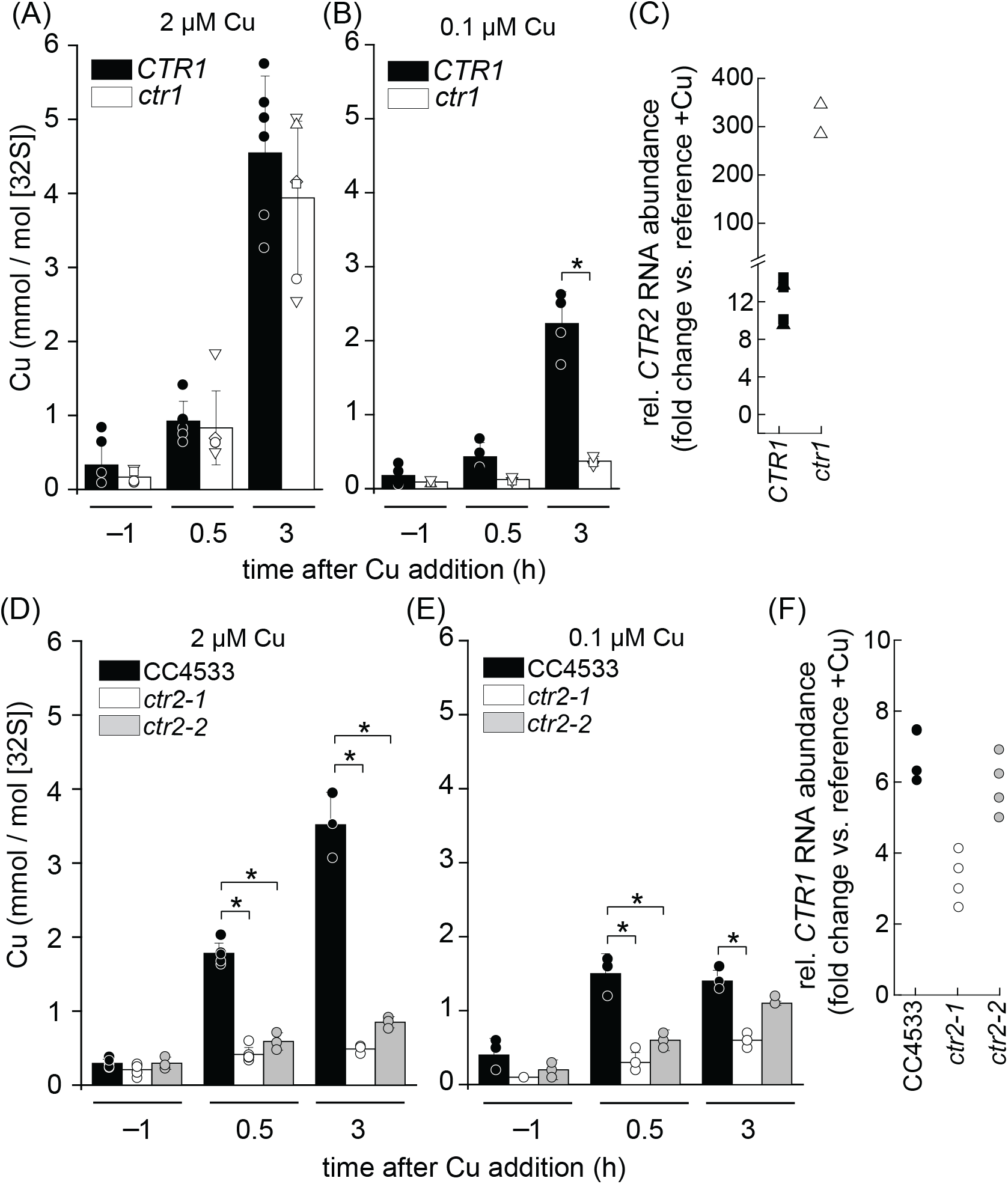
CTR1 and CTR2 are both functional Cu importers *in vivo*, albeit with distinct functional properties. (ABDE) Cu addition experiment was performed as described in Figure 1. Cu content measured by ICP-MS/MS. Individual data points are shown with averages indicated by the bars and STDEV of 3-6 independent experiments. (CF) Relative transcript abundances of *CTR1* and *CTR2* were determined using quantitative RT-PCR. Data points, each representing an independent experiment, indicate the fold-difference in Cu-deficient vs. -replete medium.

The distinct outcome from the addition of 100 nM Cu or 2 µM Cu to the *ctr1* cultures suggests that CTR1 is a high-affinity Cu transporter, while CTR2 may be important for low-affinity Cu transport. We conclude that CTR2 cannot rescue *ctr1* in low Cu despite a striking ∼ 30-fold increase in *CTR2* transcript levels in the *ctr1* mutant strains (Figure 6C); under high Cu conditions, CTR2 can compensate for the loss of CTR1. CTR1, conversely, can rescue the *ctr2* mutants in medium containing low Cu, but not high Cu, and *CTR1* transcripts are also not increased in abundance in Cu deficient *ctr2* mutants (Figure 6F). This anomaly can be explained if CTR1 does not function when Cu content in the growth medium is high. For instance, CTR1 may be removed from the plasma membrane by ubiquitylation and degradation or by endocytosis [39]. The results of the above experiments highlight the distinct molecular properties of CTR1 and CTR2 *in vivo*: CTR1 is most relevant when Cu availability is low, while CTR2 is more relevant when Cu concentrations are higher.

### CTR transporters are responsible for luxury Cu uptake in Zn-deficient cells

Zn is another element that is required at trace levels for all forms of life because of its function as an electrophile. Like Cu, it is at the top of the Irving-Williams series and binds tightly to functional groups found in proteins. Metabolic connections between Cu and Zn have been observed in some organisms, and in general the molecular basis for these connections is not known. We determined that *crr1* cells are more compromised for growth in Zn-deficient medium compared to the complemented *CRR1* strain (Supplemental Figure 10), suggesting a role for CRR1 in low Zn conditions. When we tested individual *ctr* mutants to assess whether loss of CTR function might be responsible for the *crr1* phenotype in Zn-deficient conditions, we observed that none of the *ctr* strains show a growth phenotype similar to that of *crr1* (Supplemental Figure 10) even though the abundance of *CTR2* and *CTR3* transcripts increases in Zn-deficient cells (Figure 7A). The magnitude of the increase in *CTR* expression is smaller than that in Cu-deficient cells, but it is significant and correlates with unusually high Cu content in Zn-deficient cells [40].

**Figure 7:**
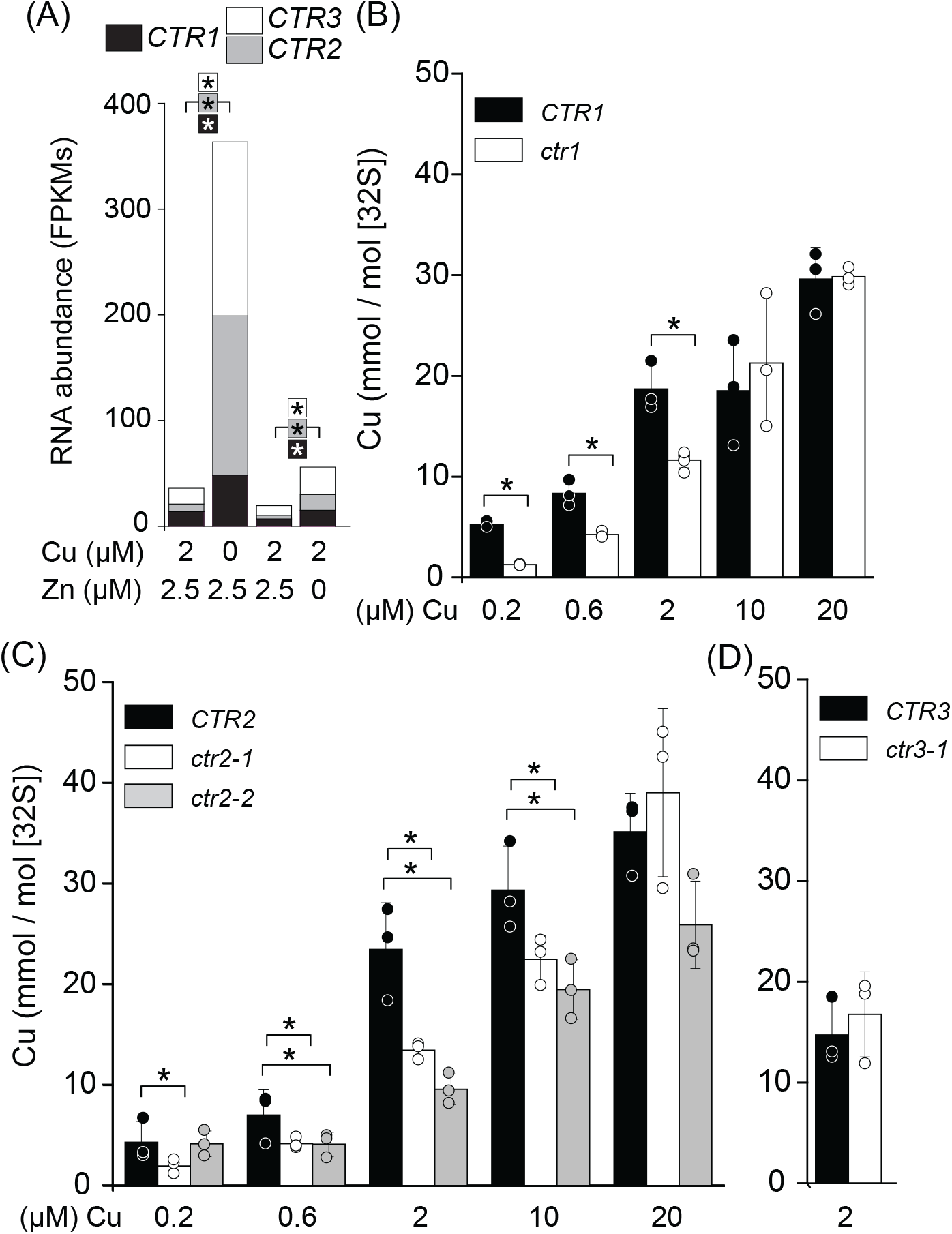
Copper accumulation is impaired in zinc deficient *ctr1* and *ctr2* mutants. (A) Expression of *CTR1*, *CTR2* and *CTR3* is increased in Zn deficiency relative to sufficiency. RNA sequencing data from (Castruita et al., 2011; Hong-Hermesdorf et al., 2014). (BCD) Cu content of *ctr1*, *ctr2* and *ctr3* mutants and corresponding reference strains grown in Zn deplete media, supplemented with Cu as indicated, was measured using ICP-MS/ MS. Shown are data points, averages and StDEV of at least three independent experiments.

Therefore, we tested each *ctr* mutant for Cu accumulation under Zn deficiency (Figure 7). The *ctr1* and *ctr2* mutants accumulated less Cu compared to the corresponding wild-type strains, specifically when Cu was supplied at a range of 0.2 µM to 2 µM in the growth medium, with *ctr1* displaying a stronger phenotype, compatible with the model that CTR1 is the higher affinity transporter. When Cu concentrations in the medium were increased to 10 µM or 20 µM, Cu accumulation in *ctr1* matched that in the wild type (Figure 7B, C). This finding raised the possibility that the same strategy might allow *crr1* mutants to grow under Zn deficiency and will allow this mutant to also overaccumulate Cu (see below).

Taken together, CTR1 and CTR2 both contribute to Cu accumulation in Zn deficiency, with CTR1 being more relevant when external Cu availability is low and CTR2 contributing more at elevated Cu concentrations.

### Nutritional complementation of the *crr1* mutant with Cu distinguishes Cu transport into and out of the acidocalcisome

We wondered if we might be able to force Cu uptake into the *crr1* mutant by growing the strain in Zn-deficient conditions. Because *crr1* grows poorly under prolonged Zn-deficient conditions, we first grew *crr1* and *CRR1* strains in medium replete for Zn and Cu and collected 2 x 10^8^ cells, followed by washing cells twice with 1 mM EDTA to remove bound metals from the cell surface before transfer to fresh Zn-deficient medium to a final cell density of 2 x 10^6^ cells/mL (comparable to a log phase culture). For the *CRR1* strain, we added 1.2 µM Cu to the medium, but chose 40 µM in the case of *crr1* to overcome any possible uptake defect from loss of expression of both *CTR1* and *CTR2* (Figure 8AB). This protocol allowed growth of the *crr1* mutant in Zn deficiency for 2 days, leading to Cu accumulation in *crr1* comparable to that in the wild-type strain at 24 h into transfer to Zn-deficient, Cu-excess conditions (Figure 8C). Thus, excess Cu in the external medium drove some Cu into *crr1* experiencing Zn deficiency, which facilitated analysis of the roles of CTR1 and CTR2 in intracellular Cu distribution.

**Figure 8:**
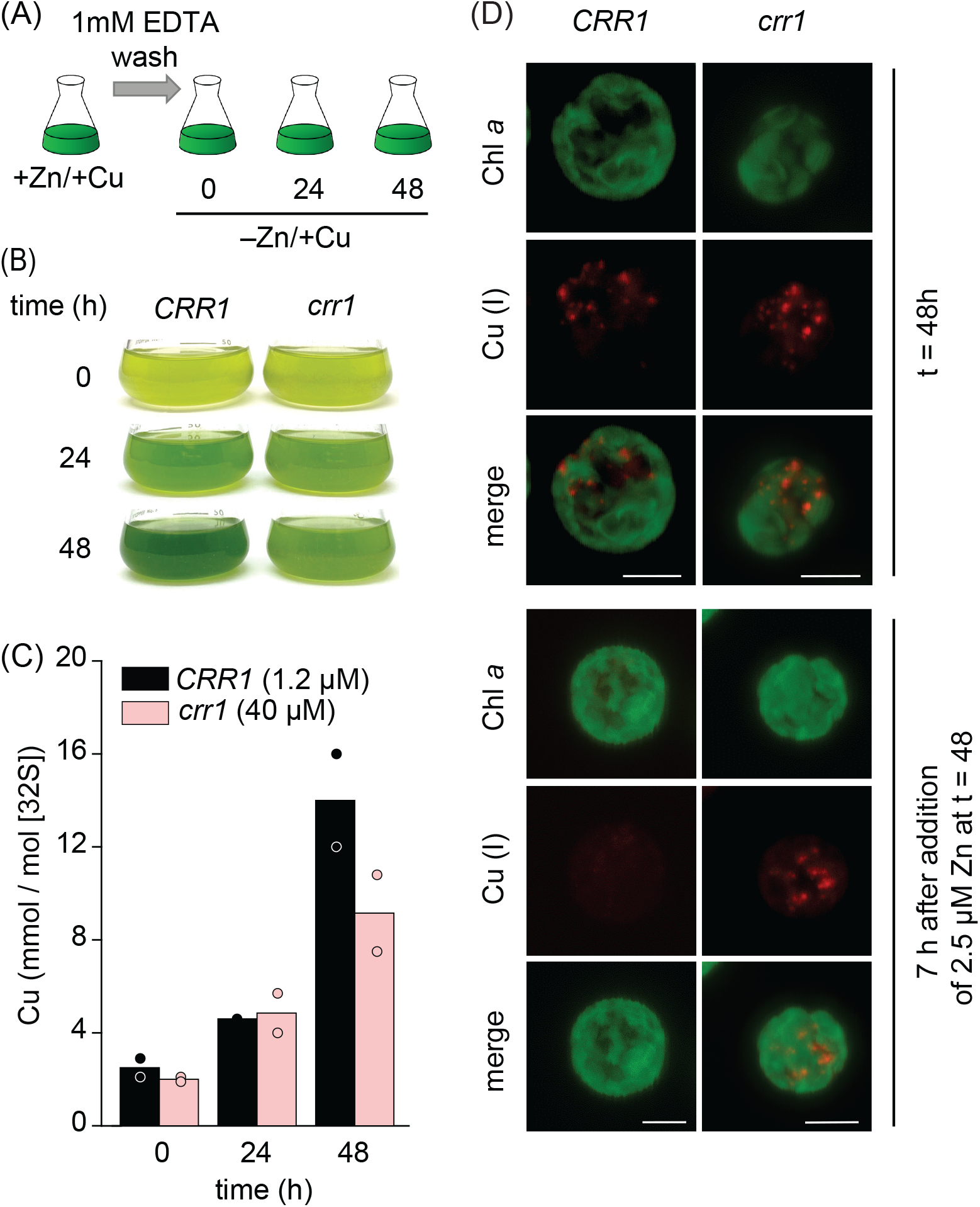
Excess Cu accumulates in Zn deficient *crr1* and is stored in acidocalcisomes, but cannot be mobilized after Zn add back. Cells were grown in copper replete medium, washed with 1mM EDTA and resuspended in Zn deplete media as shown in (A). (B) Pictures of flasks were taken every 24 h during the experiment. (C) Cu content of *CRR1* and *crr1* cell lines was measured using ICP-MS/MS. Growth medium was supplied with either 1.2µM (*CRR1*) or 40µM Cu (*crr1*) to Zn-deplete growth medium as indicated. Shown are averages and individual data points of two independent experiments. (D) After 48 h in Zn deficiency, Zn was added back to the culture medium. Samples for imaging were taken right before and 7 h after Zn add back. Cu was visualized using the Cu(I) sensitive dye CS3 (red), cells were visualized using chlorophyll autoflourescence (green). Shown are max intensity projections of each channel. Scale bars represents 5 µm. Experiment was performed at least twice with independent cultures. At least 6 cells were imaged and are shown in Supplemental Figure 11.

In wild-type cells, excess Cu localizes to the acidocalcisome, a low-pH compartment containing polyphosphate and calcium ions (Ca^2+^) [40]. When we stained cells with the Cu dye Coppersensor-3 (CS3) to visualize Cu(I) storage sites, we detected distinct foci in the *crr1* and *CRR1* strains at 48 h after transfer to Zn-deficient medium (Figure 8D). We conclude that excess external Cu can bypass the requirement for CTR transporters for Cu uptake into cells. The subsequent Cu sequestration into the acidic vacuoles occurs independently of CRR1 and CTRs. To test whether this pool of Cu can be remobilized, we supplied cultures with 2.5 µM Zn (a typical Zn-replete condition): we noticed the release of Cu from vacuoles in the *CRR1* strain within 7 h, in agreement with previous results [40], but not from the *crr1* mutant (Supplemental Figure 11). We conclude that the transporter responsible for efflux of Cu(I) from the acidocalcisome is a target of CRR1.

We investigated the possible contribution of each Cu transporter to Cu efflux from the acidocalcisome by probing each *ctr* mutant for mobilization of stored Cu(I) from the acidocalcisome. The *ctr* mutants accumulated Cu to a moderate level when grown under Zn deficiency with standard 2 µM Cu supplementation (Figure 7B, C): the accumulated Cu(I) was visible as distinct foci corresponding to the acidocalcisomes (Figure 9 upper panel), similar to wild-type strains and the *crr1* mutant. We conclude that none of the transporters is likely required for sequestration into acidocalcisomes. However, upon re-supply of Zn, only the *ctr1* and *ctr3* mutant strains were able to mobilize Cu(I) from the organelle, while Cu(I) foci remained in both *ctr2* mutants (Figure 9 lower panel and Supplemental Figures 12 and 13). We conclude that CTR2 may localize, at least in part, to the boundary membrane of the acidocalcisome and functions to release Cu(I) from intracellular stores. The concentration of Cu(I) in the acidocalcisome is expected to be high in the Cu-accumulating situation driven by Zn deficiency, which is compatible with the lower substrate affinity of CTR2 for Cu(I).

**Figure 9:**
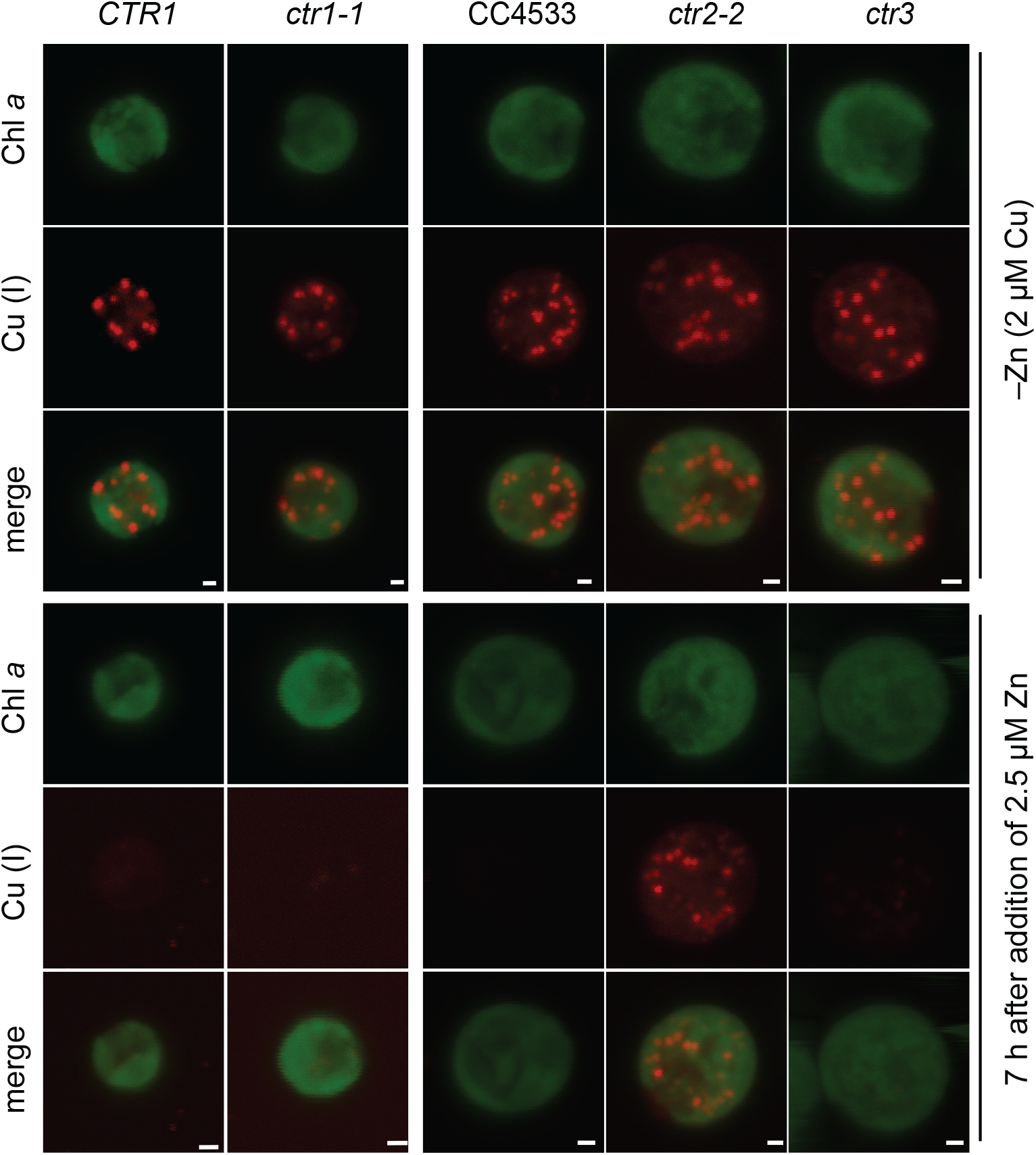
CTR2 exports Cu(I) out of the acidocalcisomes. Cells were grown in medium without Zn supplementation. At a cell density between 2-4 x 10^6^ cells/ ml, Zn was added back to the culture media. Samples for imaging were taken right before and 7 h after Zn addition. Cu was visualized using the Cu(I) sensitive dye CS3, cells were visualized using chlorophyll autofluorescence. Scale bars represents 5 µm. Experiments were performed at least twice with independent cultures. At least 5 cells of two independent mutants were imaged and are shown in Supplemental Figures 12-13.

## CONCLUSIONS

A previous study revealed that Chlamydomonas, like many other organisms, contains more than one gene encoding CTR-type Cu transporters [7]. COPT1 is most closely related to Arabidopsis Cu transporters but its expression is unchanged in response to Cu nutritional changes and its heterologous expression does not rescue the yeast *ctr1* mutant [7, 19]. In this work, we establish CTR1 and CTR2 as members of the phylogenetically distinct CTRA-CTR family and as canonical Cu transporters *in vivo*.

CTR1 and CTR2 are likely the descendants of a CTRA-CTR protein acquired from a Chytrid, which functionally replaced the laterally inherited plant-like CTR protein for Cu assimilation across the plasma membrane. Presumably, the Cu-acquisition characteristics enabled by the presence of the CTRA domains (that are shared with extant Chytrids) provided a selective advantage over the ancestral plant-like CTR protein. Another selective advantage acquired during the evolution of the Chlamydomonadales was enabled by the evolution of a secreted protein containing three CTRA domains. However, the selective advantage due to the presence of CTR3 and similar proteins has yet to be elucidated. Nevertheless, we localized CTR3 to the periplasmic space in Chlamydomonas, consistent with a predicted N-terminal signal sequence. A CTRA-containing protein, analogous to CTR3, has evolved independently in the amoeba *Dictyostelium discoideum* likely after duplication of the gene encoding the CTRA-CTR protein referred to as P80 [41]. However, whereas CTR3 has three CTRA domains, the soluble P80-derived protein has a single CTRA domain. P80 and orthologous CTRA-CTR proteins from Amoebozoa are likely also the descendants of a horizontally transferred gene from an ancient Cytrid, but the event was independent from the event that resulted in the green algal CTRA-CTR proteins.

Conservation of the putative Cu-binding ligands in the fungal, amoeba, and algal CTRA-CTR and CTRA proteins suggests that the CTRA proteins play a role in Cu homeostasis, possibly in Cu acquisition prior to transport by CTRA-CTR. However, under the conditions tested here, we did not observe a Cu-assimilation defect in the CTR3 mutant. Based on the presence of characterized and uncharacterized proteins in the CRR1 regulon, *C. reinhardtii* has acquired an arsenal to compete with other organisms for Cu and thrive during Cu limitation. As such, there may be several functionally overlapping pathways that can complement the loss of CTR3. One such pathway may involve Cre05.g236039, an unrelated, soluble protein predicted to be secreted, that contains numerous Cys, Met, and His residues. Like CTR3, orthologs of Cre05.g236039 are often fused to the N-terminus of CTR proteins in other algae and protists. The reason why some green algae employ the CTRA-CTRs, while others employ the Cre05.g236039-like-CTR fusions is unknown, but the presence of CTR3, Cre05.g236039 and possibly other putative, extracellular, Cu acquisition proteins points to the presence of a complex, seemingly redundant Cu-acquisition strategy. In the case of Cu transport across the plasma membrane, our results presented here highlight the distinct functional roles of the paralogs CTR1 and CTR2. Retaining horizontally transferred genes and duplication events giving rise to multiple CTR paralogs with different functional properties has likely enabled Chlamydomonas and other organisms to fine-tune Cu uptake and distribution. In particular, differences in domain organization, even relatively minor differences in Cu-binding motifs, allow for functional flexibility in Cu assimilation. CTR1 functions as a high-affinity importer whose activity is turned off at high extracellular Cu to prevent Cu overload during transition of cells from very low to high Cu. This could occur by ubiquitylation followed by endocytosis or degradation. CTR2 has two functions: it is a lower affinity transporter that delivers Cu(I) to the cytoplasm from intracellular stores where the Cu(I) content is high and also enables Cu(I) uptake but only at high Cu content in the medium. The combination of transporters with different affinities, localization and trafficking allows tight control of cytoplasmic Cu availability for cuproprotein synthesis.

## METHODS

### Generation of ctr1 KO strains using CRISPR/Cpf1

CC-425, a cell wall reduced arginine auxotrophic strain, was used for transformation with a ribonucleoprotein (RNP) complex consisting of a guide RNA (gRNA) targeting a PAM sequence in exon1 of *CTR1* and LbCpf1 as described in [35, 37]. Modifications are outlined as follows: cultures were grown to 2 × 10^6^ cells per mL and counted by using a hemocytometer. For optional pretreatment, 2 × 10^7^ cells were suspended and centrifuged (5 min, 1424 x g) in Maxx Efficiency Transformation Reagent (1 mL) twice, followed by suspension in 230 μL of the same reagent supplemented with sucrose (40 mM). Cells were incubated at 40°C for 20 minutes. Purified LbCPF1 (80 μM) was preincubated with gRNA (1 nmol, targeting sequence = TTTGGGATGCGGCGGCGCTCAGCGG) at 25 °C for 20 min to form RNP complexes. For transfection, 230 μL cell culture (5 × 10^5^ cells) was supplemented with sucrose (40 mM) and mixed with preincubated RNPs and *Hin*DIII digested pMS666 containing the *ARG7* gene conferring the ability to grow without arginine. In order to achieve template DNA-mediated editing, an ssODN containing two stop codons after the PAM target site (∼4 nmol, sequence = GTAGCTACTGACGTGTGCAGCTCGTTCATTTGGTAAG-CGTAGGCGCTCAGCGGCTACTGCACGACGACCGGCGCTCTGGCGTACCGGTCG) was ad-ded. The final volume was 280 μL. Cells were electroporated in a 4-mm gap cuvette (Biorad) at 600 V, 50 μF, 200 Ω using a Gene Pulser Xcell (Bio-Rad). Immediately after electroporation, 800 μL of TAP with 40 mM sucrose was added. Cells were recovered overnight in darkness and without shaking in 5 mL TAP with 40 mM sucrose and polyethylene glycol 8000 0.4% (w/v) and then plated on TAP after a 5 min centrifugation at RT and 1424 x *g* using the starch embedding method. Since we did not expect a phenotype, colonies were screened by qPCR for successful genome editing at the *CTR1* locus. Colony qPCR of transformants was performed as follows: after one week of growth, half of each colony on TAP plates was resuspended in 50 uL 10 mM Tris-HCl (pH 8) buffer in a 96 well plate. Cells were heated to 96 °C for 10 min. After vortexing, 96 well plates were spun down for 4 min at ∼1000 x *g*. Supernatants containing the DNA were transferred to new 96 well plates. qPCR on a total of 178 clones was performed using oligos Ctr1screenfor CAGCTCGTTCATTTGGGATG (which was specific for the wild-type, unedited *CTR1* gene) and Ctr1unirev GTGTGAGAGCTGGCTGATCC. The following program was used for all qPCR reactions: 95°C for 5 min followed by 40 cycles of 95°C for 15 s, and 65°C for 60 s. 11 clones failed to amplify the wild-type CTR1 sequence; these were subjected to a secondary screen using oligo Ctr1seqfor ACGTGTGCAGCTCGTTCATT (specific for gene-edited *ctr1* alleles) and Ctr1unirev GTGTGAGAGCTGGCTGATCC. Sequencing of the PCR products using oligo Ctr1unirev revealed that 5 clones had ssODN-mediated gene editing within the first exon of CTR1, and all of them showed successful introduction of the two stop codons. *ctr1-3* also contained three extra base pairs integrated downstream of the gene editing target site in addition to the two stop codons in exon1 (Supplemental Figure 1).

### Culture conditions and additional Chlamydomonas strains

Insertional mutant lines from the Chlamydomonas Library Project (CliP) [38] were designated as *ctr2-1* (LMJ.RY0402.151308), *ctr2-2* (LMJ.RY0402.163662), *ctr3-1* (LMJ.RY0402.179604). The mutants and corresponding wildtype strain CC-4533 were genotyped by PCR and amplicons were sequenced to confirm predicted integration of the insertion cassette. The *crr1* mutants (CC5068) and *crr1* lines complemented with *CRR1* genomic sequence (CC-5070, designated as *CRR1*) were characterized previously [42, 43]. Unless stated otherwise, cells were grown in Trisacetate-phosphate (TAP) with constant agitation in an Innova incubator (160 rpm, New Brunswick Scientific, Edison, NJ) at 24°C in continuous light (90 µmol m-2 s-1), provided by cool white fluorescent bulbs (4100K) and warm white fluorescent bulbs (3000 K) in the ratio of 2:1. Cell wall reduced strains CC-425 and CC-5390 were grown under the same light regime but with constant agitation at 140 rpm. TAP medium with or without Cu or Zn was used with revised trace elements (Special K) instead of Hunter’s trace elements [44].

### Precipitation of secreted proteins

Cells from strain CC-5390 were analyzed after three rounds of growth in Cu-deficient TAP medium to stationary phase. Cells were collected by centrifugation at 4°C for 10 min at 1424 x *g*. Following centrifugation, pellets were resuspended in a lysis solution (125 mM Tris-HCl pH 6.8, 20% glycerol, 4% SDS, 10% β-mercaptoethanol, 0.005% bromophenol blue). After a second centrifugation at 4°C for 10 min at 4000 rpm, the supernatant was filtered through a 0.4-µm filter before ice-cold 100% trichloroacetic acid (TCA) was added to the samples to a final concentration of 10%. Samples were mixed and incubated overnight at 4°C. The samples were then centrifuged at 4°C for 10 min at 1424 x *g* and resuspended in ice-cold acetone. Another centrifugation at 4°C for 10 min at 1424 x *g* was performed before final protein pellets were resuspended in 2 x sample buffer (125 mM Tris-HCl pH 6.8, 20% [v/v] glycerol, 4% SDS, 10% β-mercaptoethanol, 0.005% bromophenol blue) and incubated for 10 min at 65°C before storage at –80°C.

### Antibody production and protein analyses by immunodetection

Antibodies against CTR1 were produced by COVANCE via immunization using the subcutaneous implant procedure of rabbits following a 118-day protocol with synthetic peptide Ac-CNAKAR-RGSGDALGANTADHKKGASS-amide. For analysis of total proteins, 15 mL of a Chlamydomonas culture with a cell density of 4-8 x 10^6^ cells/mL was centrifuged for 3 mins at 4°C and at 1650 x *g*. The resulting cell pellet was resuspended in 300 µL of a buffer containing 10 mM Na-phosphate buffer (pH 7.0), EDTA-free Protease inhibitor (Roche), 2% (w/v) SDS, and 10% (w/v) sucrose. For analysis of soluble and membrane fractions, 15 mL of a culture at a cell density of 4-8 x 10^6^ cells/ mL was centrifuged at 1650 ×*g*. The cell pellet was resuspended in 300 µL of buffer containing 10 mM Na-phosphate buffer (pH 7.0) and EDTA-free Protease inhibitor (Roche). Soluble and membrane fractions were isolated by lysing cells with three freeze-thaw cycles (−20°C to room temperature). After lysis, soluble proteins (supernatant) were separated from the membrane bound proteins (pellet) by centrifugation at 4°C. Membrane-bound proteins were resuspended in 200 µL Na-phosphate buffer containing 2% Triton X-100 before both fractions were quick-frozen in liquid N_2_. Samples were stored at −80°C prior to analysis. Protein amounts were determined using a Pierce BCA Protein Assay Kit against BSA as a standard and diluted with 2 x sample buffer (125 mM Tris-HCl pH 6.8, 20% [v/v] glycerol, 4% [w/v] SDS, 10% [v/v] β-mercaptoethanol, 0.005% [w/v] bromophenol blue). Proteins were separated on SDS-containing polyacrylamide gels using 10 μg of protein per lane. The separated proteins were then transferred by semi-dry electroblotting to nitrocellulose membranes (Amersham Protran 0.1 NC). The membrane was blocked for 30 min with 3% (w/v) dry non-fat milk in phosphate buffered saline (PBS) containing 0.1% Tween 20 and then incubated in primary antibody at room temperature. PBS was used to dilute both primary and secondary antibodies. The membranes were washed in PBS containing 0.1% (v/v) Tween 20. Antibodies directed against CF_1_ (1:40,000), OEE1 (1:8,000), plastocyanin/Cyt *c*_6_ (1:4,000), CTR3 (1:1,000) and CTR2 (1:1,000) (previously used in [7], CTR1 (1:1,000), FEA1/2 (1:20,000). The secondary antibody used was a goat anti-rabbit antibody (1:5,000 dilution) conjugated to alkaline phosphatase, and processed according to the manufacturer’s instructions.

### Quantitative elemental analysis

Cell cultures corresponding to 1 × 10^8^ cells (culture density of 3–5 × 10^6^ cells/mL) were collected by centrifugation at 1424 x *g* for 3 min in a 50-mL falcon tube. The cells were washed twice in 50 mL of 1 mM Na_2_EDTA pH 8.0 (to remove cell surface–associated metals) and once in Milli-Q water. The cell pellet was stored at −20°C before being overlaid with 286 µL 70% (v/v) nitric acid and digested at room temperature for 24 h and 65°C for about 2 h before being diluted to a final nitric acid concentration of 2% (v/v) with Milli-Q water. Complementary aliquots of fresh or spent culture medium were treated with nitric acid and brought to a final concentration of 2% (v/v) nitric acid. Metal, sulfur and phosphorus contents were determined by inductively coupled plasma mass spectrometry (ICP-MS/MS) on an Agilent 8800 Triple Quadropole ICP-MS instrument by comparison to an environmental calibration standard (Agilent 5183-4688), a sulfur (Inorganic Ventures CGS1) and phosphorus (Inorganic Ventures CGP1) standard. ^89^Y served as an internal standard (Inorganic Ventures MSY-100PPM). The levels of analytes were determined in MS/ MS mode. ^23^Na, ^24^Mg, ^31^P, ^55^Mn, ^63^Cu and ^66^Zn analytes were measured directly using He in a collision reaction cell. ^39^K, ^40^Ca and ^56^Fe were directly determined using H^2^ as a cell gas. ^32^S was determined via mass-shift from 32 to 48 utilizing O_2_ as a cell gas. An average of four technical replicate measurements was used for each individual biological sample. The average variation between technical replicate measurements was 1.1% for all analytes and never exceeded 5% for an individual sample. Triplicate samples (from independent cultures) were also used to determine the variation between cultures. Averages and standard deviations between these replicates are shown in the figures.

### Confocal microscopy using CS3 dye

CS3 dye was synthesized as described in Supplemental File 1. Cell-walled Chlamydomonas cells were cultured to early stationary phase and 1–2 x 10^7^ cells very collected by centrifugation at room temperature at 3,500 x *g*, for 2 min. The supernatant was discarded and the cell pellet was resuspended in 10 mM Na-phosphate buffer (pH 7.0) containing 10 µM CS3 dye. To avoid mechanical stress to cell wall-reduced cells, they were not centrifuged, but 10 µM CS3 dye was added directly to an aliquot of the culture instead. Microscopy was performed on a Zeiss LSM880. The following excitations and emissions were used: CS3 ex/em 514 nm/537 nm, chlorophyll 633 nm/696 nm. All aspects of image capture were controlled via Zeiss ZEN Black software, including fluorescent emission signals from probes and/or chlorophyll.

### Statistical Analyses

Unless stated otherwise, a one-way ANOVA was used to test for differences between samples. Significance indicated by ANOVA was followed by a Holm-Sidak post-hoc test. Asterisks indicates a p-value of <0.05.

### Bioinformatic analyses

The sequence similarity network (SSN) of the CTR family was constructed using the EFI-EST webtool (http://efi.igb.illinois.edu/efi-est/) [45] with an alignment score of 15; nodes were collapsed based on 70% sequence identity. Sequences for inclusion in the network were identified based on the presence of the CTR domain defined by IPR007274 from the InterPro database [46] or identified in a jackhammer search [47] using CTR3 as the query. The network was visualized with Cytoscape v3.10.1 using the Prefuse Force Directed OpenCL Layout. For the phylogenetic reconstruction, CTR1 and CTR3 protein sequences were used to search the UniProt database [48] with blastp. After manually filtering low-quality hits, 285 protein sequences were used to build an approximately maximum-likelihood phylogenetic tree with MAFFT [49] and FastTreeMP [50] and on the CIPRES Science Gateway [51] with default parameters. The resulting tree was visualized and annotated in iTol [52]. Branches with less than 0.5 bootstrap support were deleted. Precomputed AlphaFold predictions [53] were downloaded from the AlphaFold Protein Structure Database (https://alphafold.ebi.ac.uk/). Sequence logos were visualized with Skylign [54].

## ACKNOWLEDGMENTS

The work was supported by a grant from the National Institutes of Health GM42143. Work at the Molecular Foundry was supported by the Office of Science, Office of Basic Energy Sciences, of the U.S. Department of Energy under Contract No. DE-AC02-05CH11231. Work at the U.S. Department of Energy Joint Genome Institute (https://ror.org/04xm1d337), a DOE Office of Science User Facility, is supported by the Office of Science of the U.S. Department of Energy operated under Contract No. DE-AC02-05CH11231. Confocal microscopy experiments were conducted using a Zeiss LSM 880, at the CRL Molecular Imaging Center, supported by the Helen Wills Neuroscience Institute. We would like to thank Holly Aaron and Feather Ives for their microscopy training and assistance (RRID:SCR_017852), and Chris Jeans and the QB3 Macrolab at UC Berkeley for purification of LbCpf1.

## AUTHOR CONTRIBUTIONS

Designed experiments: DS, SRS, Designed CS3 synthesis HN; Performed experiments: DS, SRS, SP, BCB, SG, CS, PAS, JLM; Analyzed data: DS, SRS, SP, SSM; Prepared figures: DS, SRS, SP, CBH; Designed project: DS, SSM; Secured funding: SSM. Wrote manuscript: DS, SSM.

## DATA AVAILABILITY STATEMENT

The data underlying this article are available in the article and in its online supplementary material.

