## Supplemental Figures and legends for "Distinct function of Chlamydomonas CTRA-CTR transporters in Cu assimilation and intracellular mobilization"

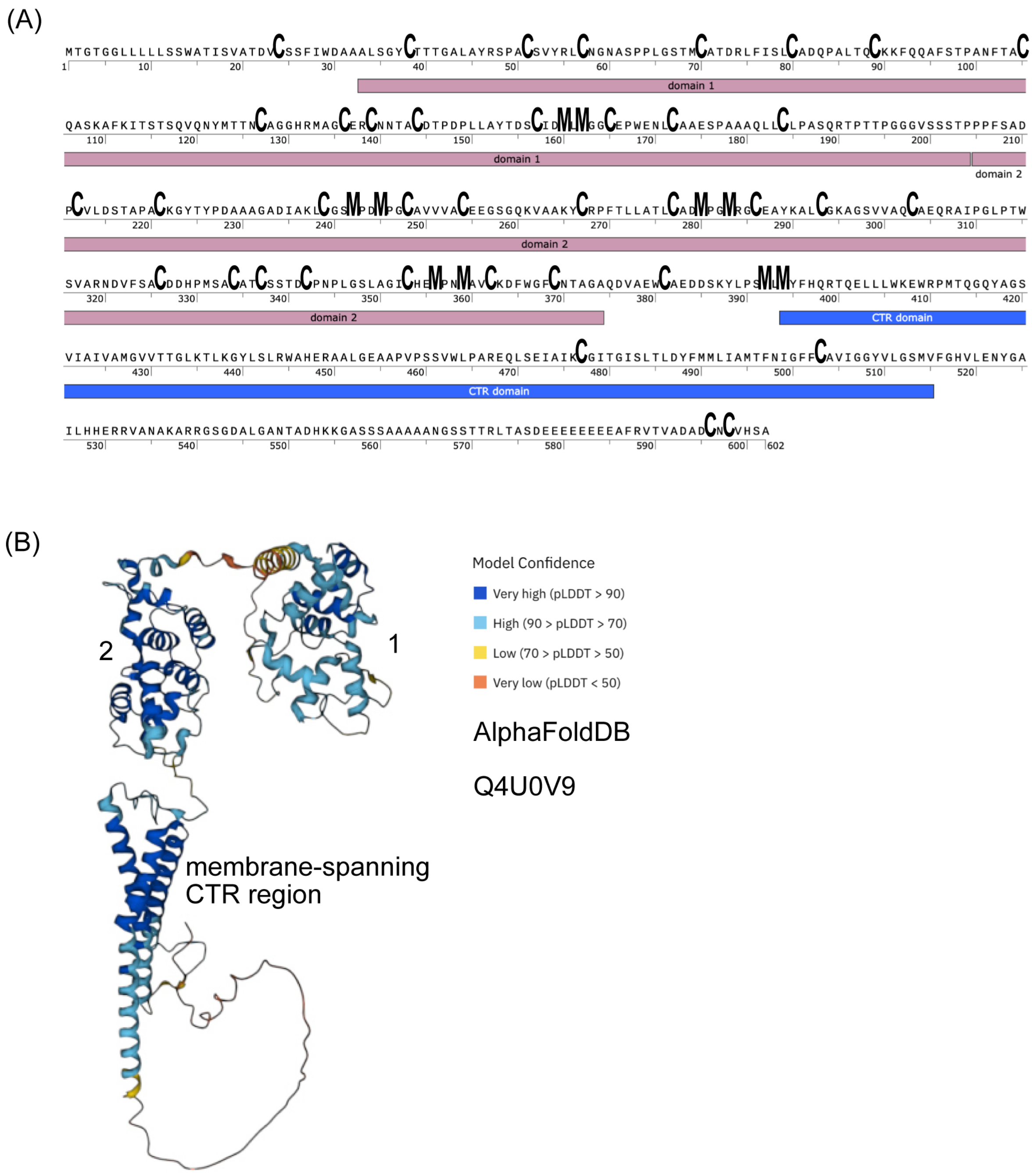


**Supplemental Figure 1.** CTR1. (A) Amino acid sequence of CTR1. Cysteine residues and Mets motifs are highlighted, as is the location of the CTRA and CTR domains. (B) AlphaFold prediction of CTR1. The three CTRA domains are numbered.

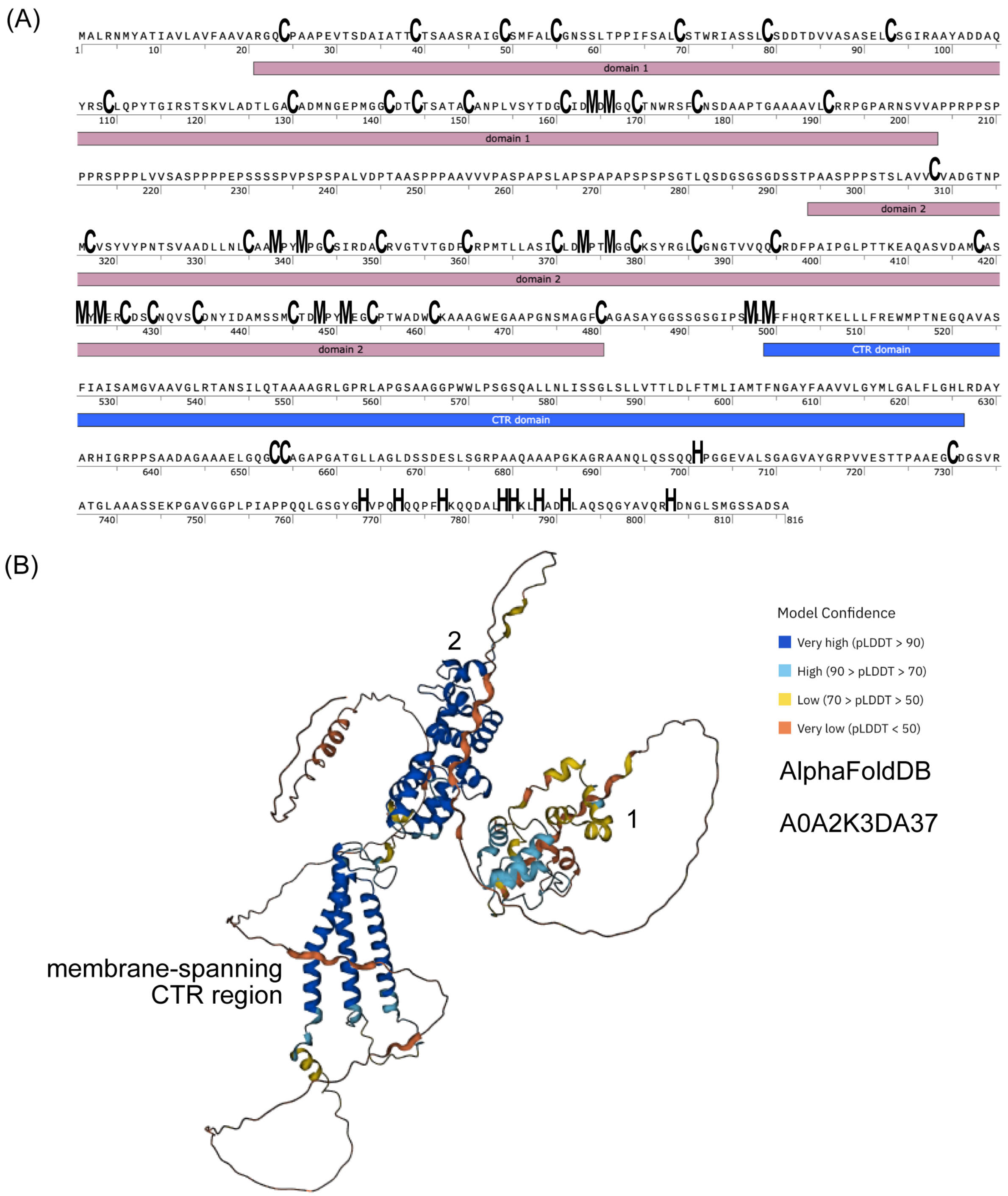


**Supplemental Figure 2**. CTR2. (A) Amino acid sequence of CTR2. Cysteine residues and Mets motifs are highlighted, as is the location of the CTRA and CTR domains, and the His-rich C-term region. (B) AlphaFold prediction of CTR2. The three CTRA domains are numbered.

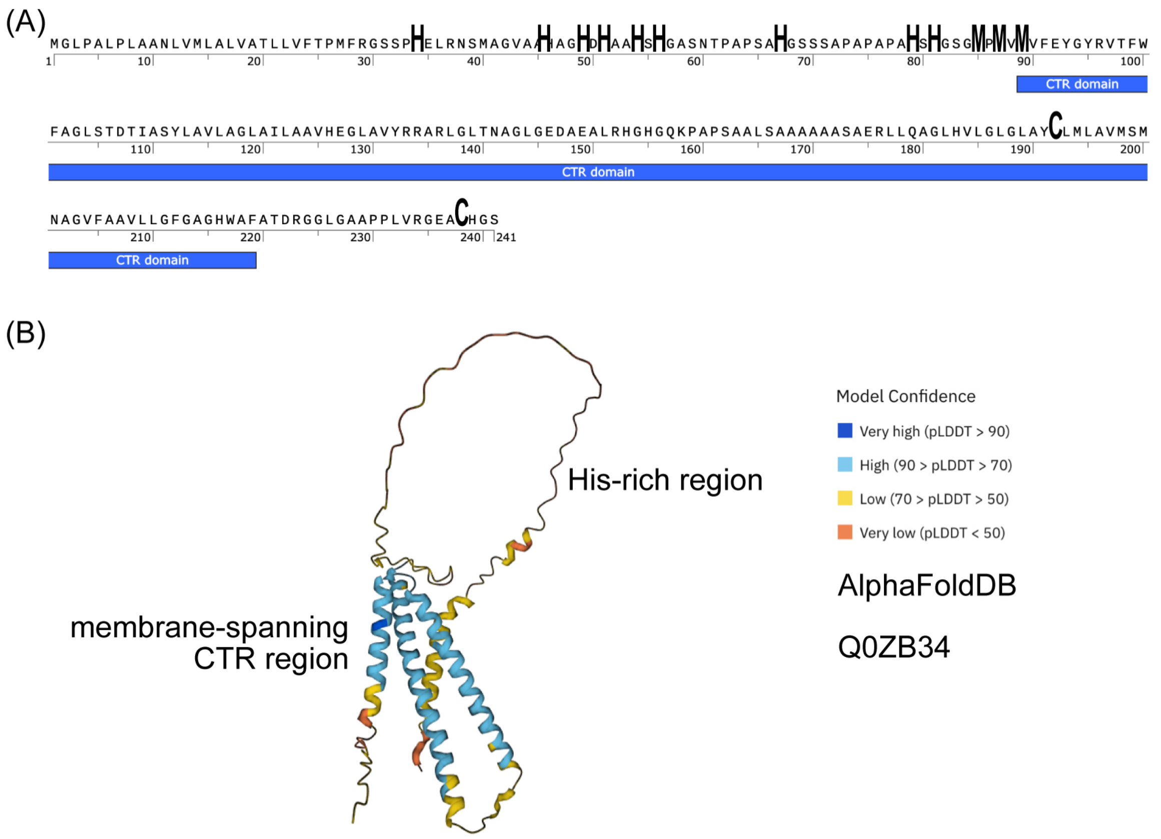


**Supplemental Figure 3**. COPT1. (A) Amino acid sequence of COPT1. Histidine residues and Mets motifs are highlighted, as is the location of the CTR domain. (B) AlphaFold prediction of COPT1.

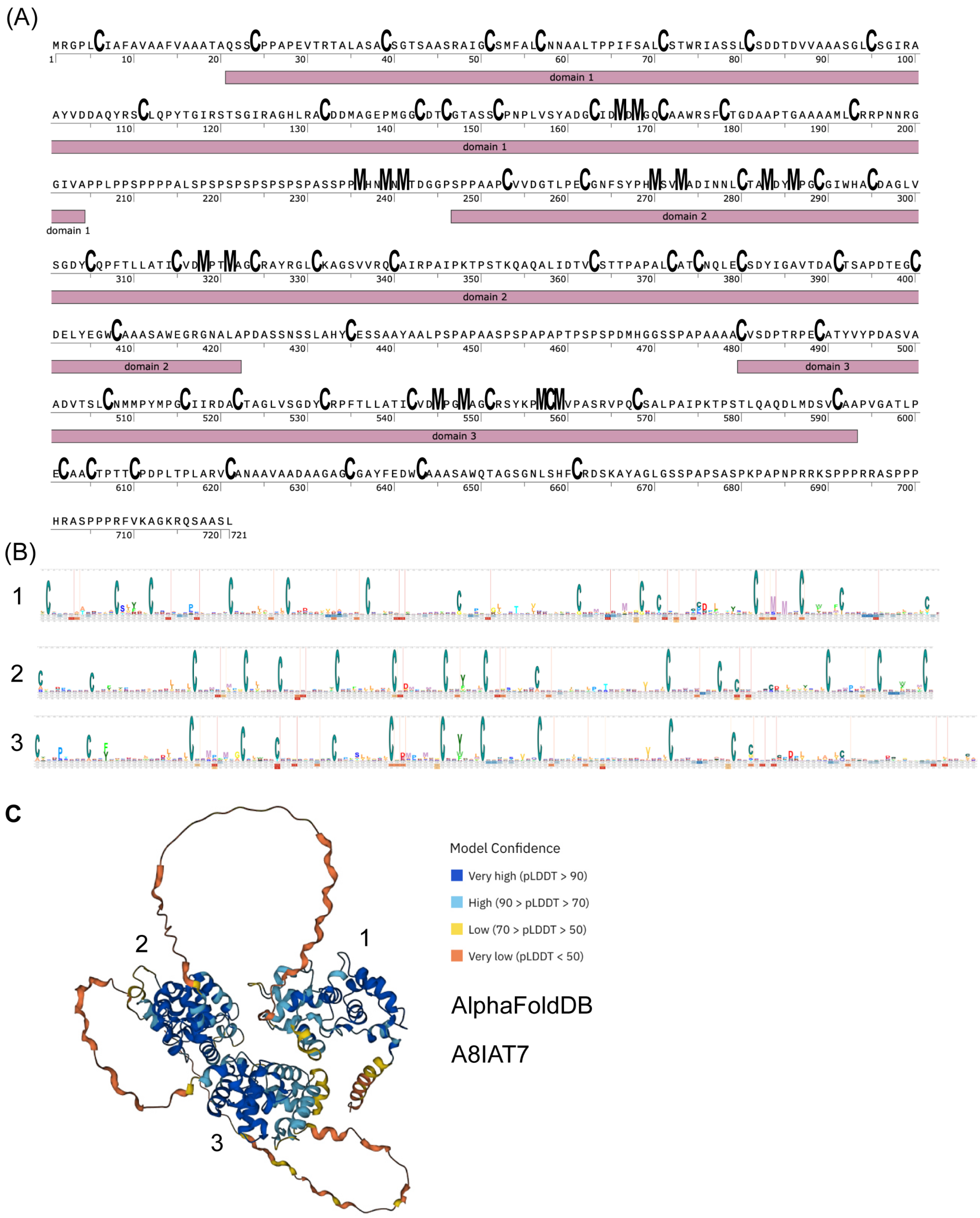


**Supplemental Figure 4.** CTR3. (A) Amino acid sequence of CTR3. Cysteine residues and Mets motifs are highlighted, as is the location of the CTRA domains. (B) AlphaFold prediction of CTR3. The three CTRA domains are numbered.

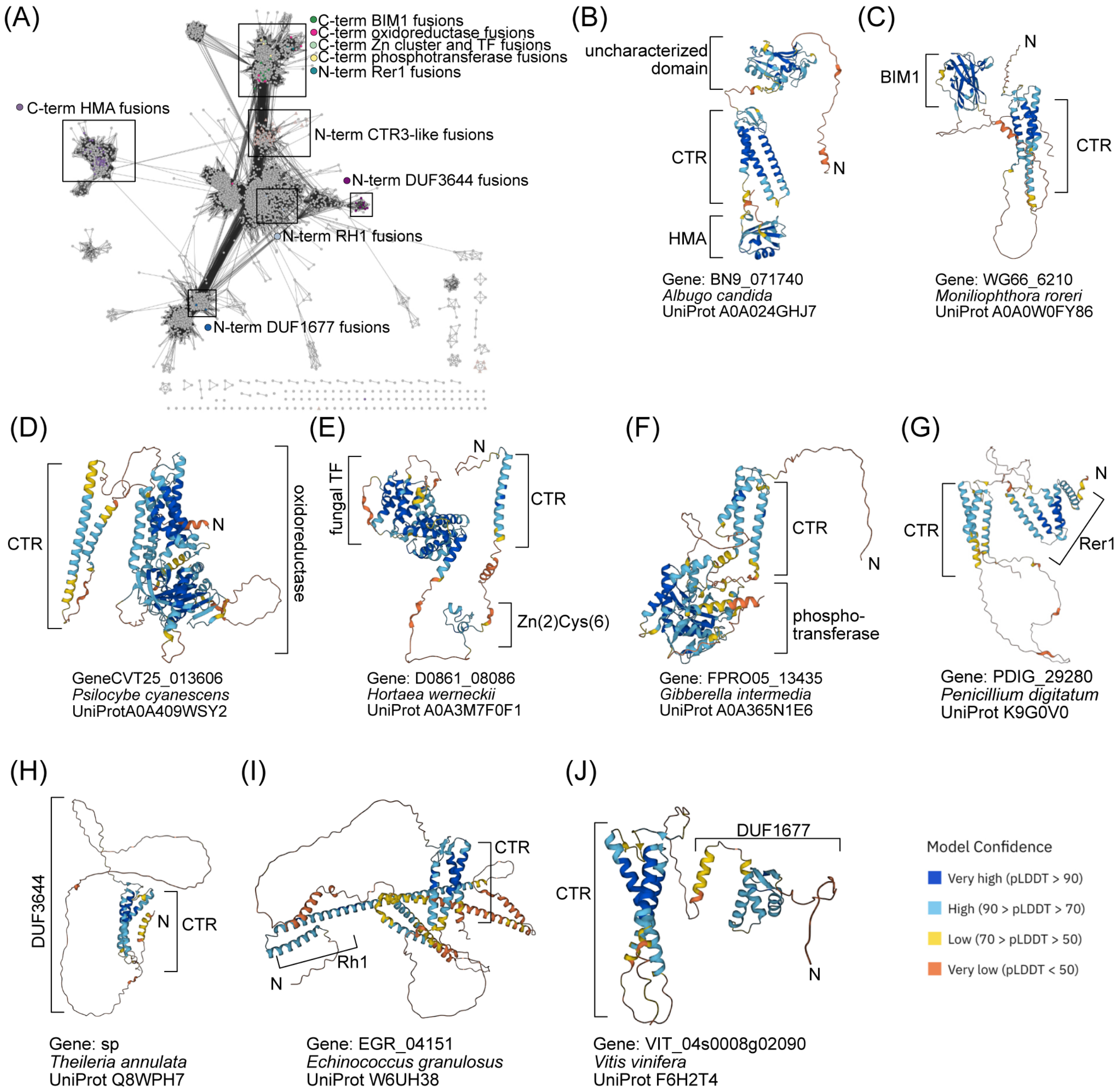


**Supplemental Figure 5.** CTR domains. (A) Sequence similarity network from Figure 2. Nodes are colored according the presence of the indicated N- or C-terminal domains. Black outlines are used to indicate the location of those fusion proteins. Only fusions that are found in at least three distinct species, and therefore less likely to be the result of inaccurate gene models, are listed. (B-J) AlphaFold prediction of representative CTR fusions shown in panel A.

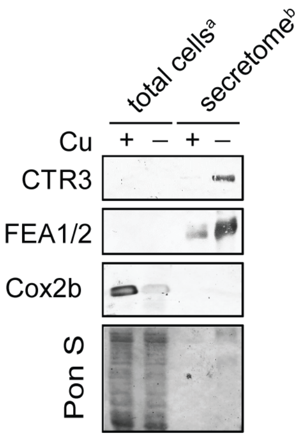


**Supplemental Figure 6: CTR3 is a secreted protein.** Total cellular protein or proteins precipitated from spent medium of a cell wall-less mutant (secretome) from Cu-deficient (–) and Cu-replete (+) media were separated by SDS-PAGE using protein derived from either [a]$2x10^6$ cells or [b]$2x10^7$ cells as indicated. Membranes were decorated with antibodies against CTR3, FEAs and Cox2b, Pon S= Ponceau stain. Shown are representative data from at least two independent experiments.

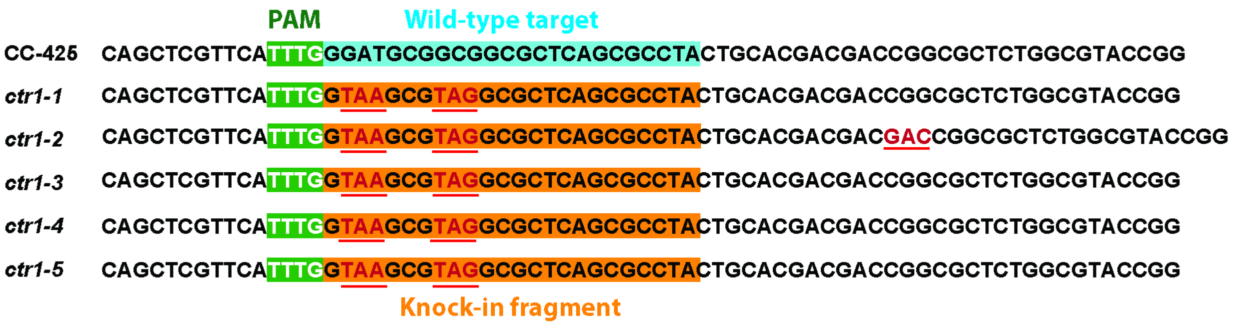


**Supplemental Figure 7. ssODN mediated CRISPR/Cpf1 gene editing introducing two in-frame stop codons in exon 1 of *CTR1*.** Sequencing of cells edited at *CTR1* using LbCpf1 (Ribonucleoproteins) RNPs. *CTR1* was amplified from arginine- resistant colonies growing on solid TAP growth medium. Green background indicates the PAM target site, blue background indicates the wild-type sequence, orange background indicates the region after the cleavage site that contains nucleic acids that deviated in the ssODN from the wild-type sequence. Stop codons are underlined in red, other deviations from the wild-type sequence are highlighted in red.

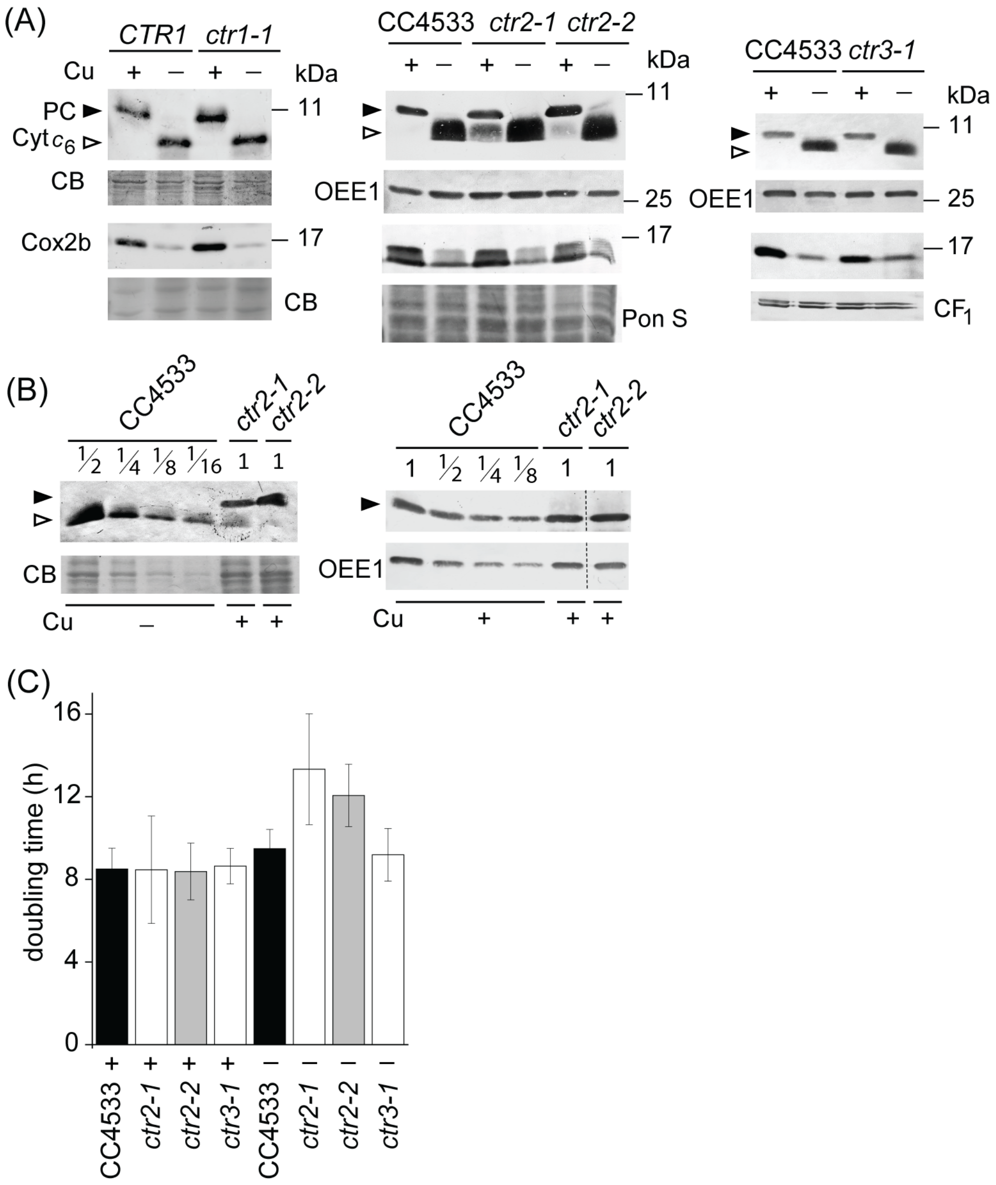


**Supplemental Figure 8: Cytochrome $c_6$ accumulation indicates internal Cu deficiency in *ctr2.* Growth is unaffected in *ctr2* and *ctr3* mutants. Cytochrome $c_6$ accumulation indicates internal Cu deficiency in *ctr2*.** *ctr1-1, ctr2-1, ctr2-2, ctr3-1* and respective reference lines were grown photoheterotrophically under Cu-deficient (–) or Cu-replete (+) conditions. (A) Abundance of Plastocyanin and cytochrome $c_6$ was determined by separating 10 µg of total soluble cell lysate using SDS-PAGE (15% monomer), followed by immunoblotting using antisera cross-reactive to plastocyanin (PC, black arrow) and Cyt $c_6$ (white arrow). In order to check abundance of Cox2b, proteins in total cell lysates were separated by SDS-PAGE (15% monomer) followed by immunoblotting for CoxIIb. Either OEE1, alpha and beta subunits of $CF_1$, Ponceau S stain (Pon S) or Coomassie blue (CB) were used as loading controls. (B) Dilution series of (left panel) Cu deficient CC4533 to determine Cyt $c_6$ abundance and of (right panel) Cu replete CC4533 to determine plastocyanin abundance in *ctr2-1* and *ctr2-2*. Shown is one example from at least two independent experiments. (C) If growth occured, cells were counted using a hemocytometer and doubling times estimated based on cell counts during the exponential growth phase. Shown are averages and StDEV of at least 3 independent experiments.

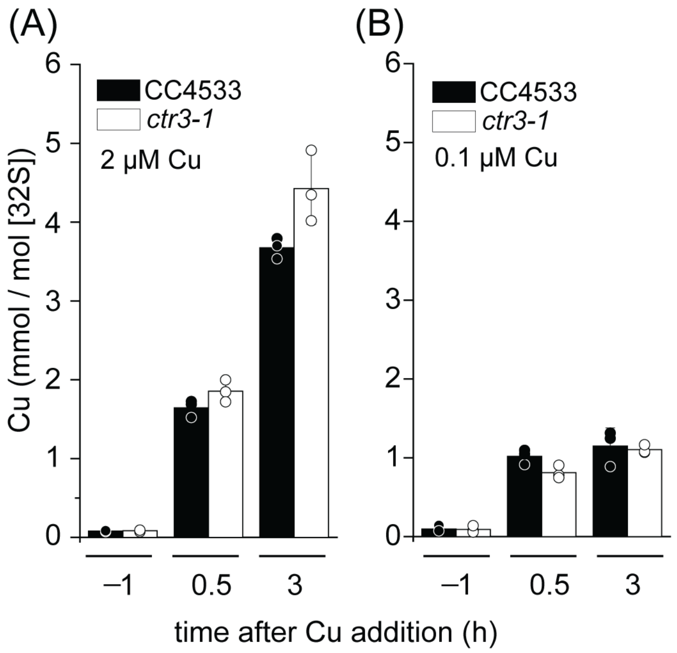


**Supplemental Figure 9:** (AB) *CTR3* and *ctr3-1* cell lines following the resupply of either 2 μM or 100 nM Cu as indicated to Cu-deficient cultures. Shown are data points, averages and StDEV of 3-6 independent experiments

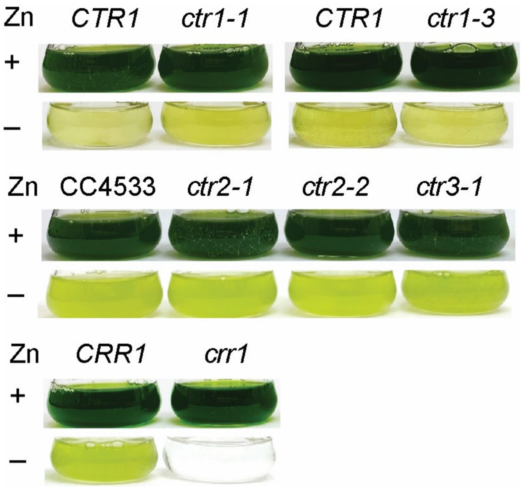


**Supplemental Figure 10. Zinc deficient growth defect is not exacerbated in *ctr* mutants.** Pictures of flasks of *ctr1*, *ctr2, ctr3* and *crr1* mutants and corresponding reference strains grown in zinc deplete media were taken eight days post-inoculation.

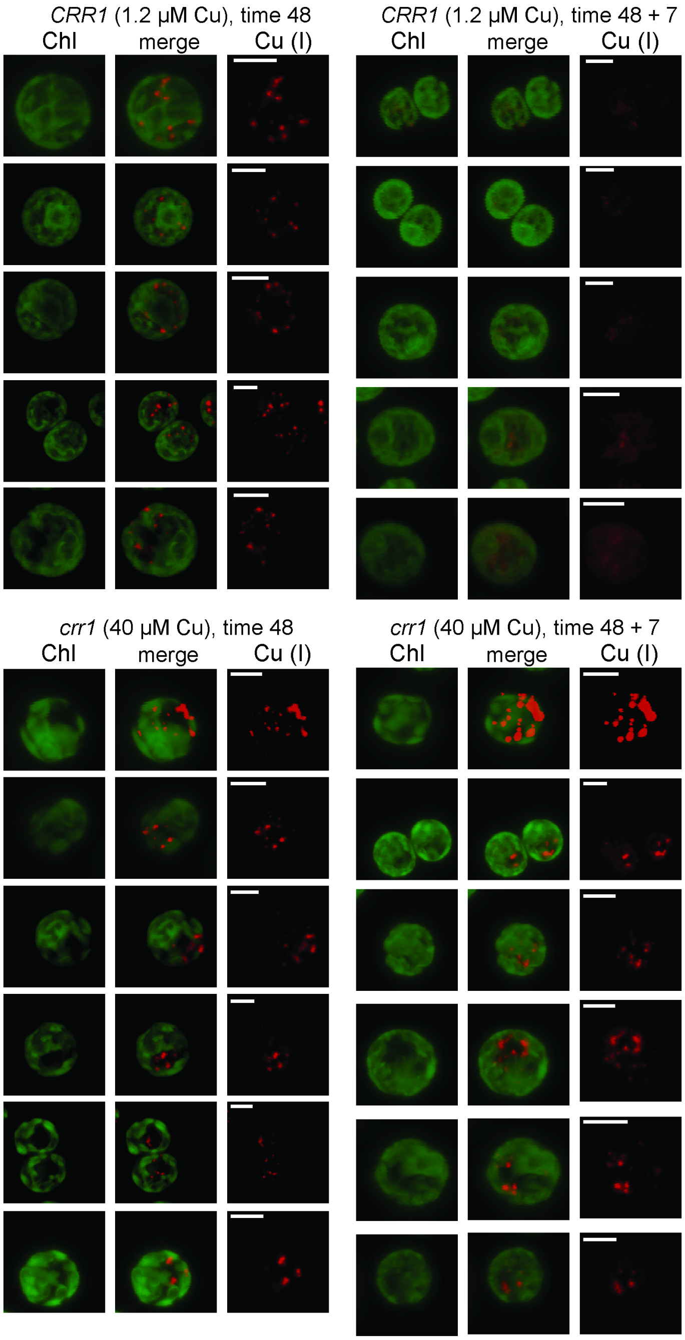


**Supplemental Figure 11. Copper accumulated in zinc deficient *crr1* is stored in distinct foci, but copper cannot be mobilized after zinc add back**. Figure shows all cells that were imaged in experiments shown and described in Figure 8.

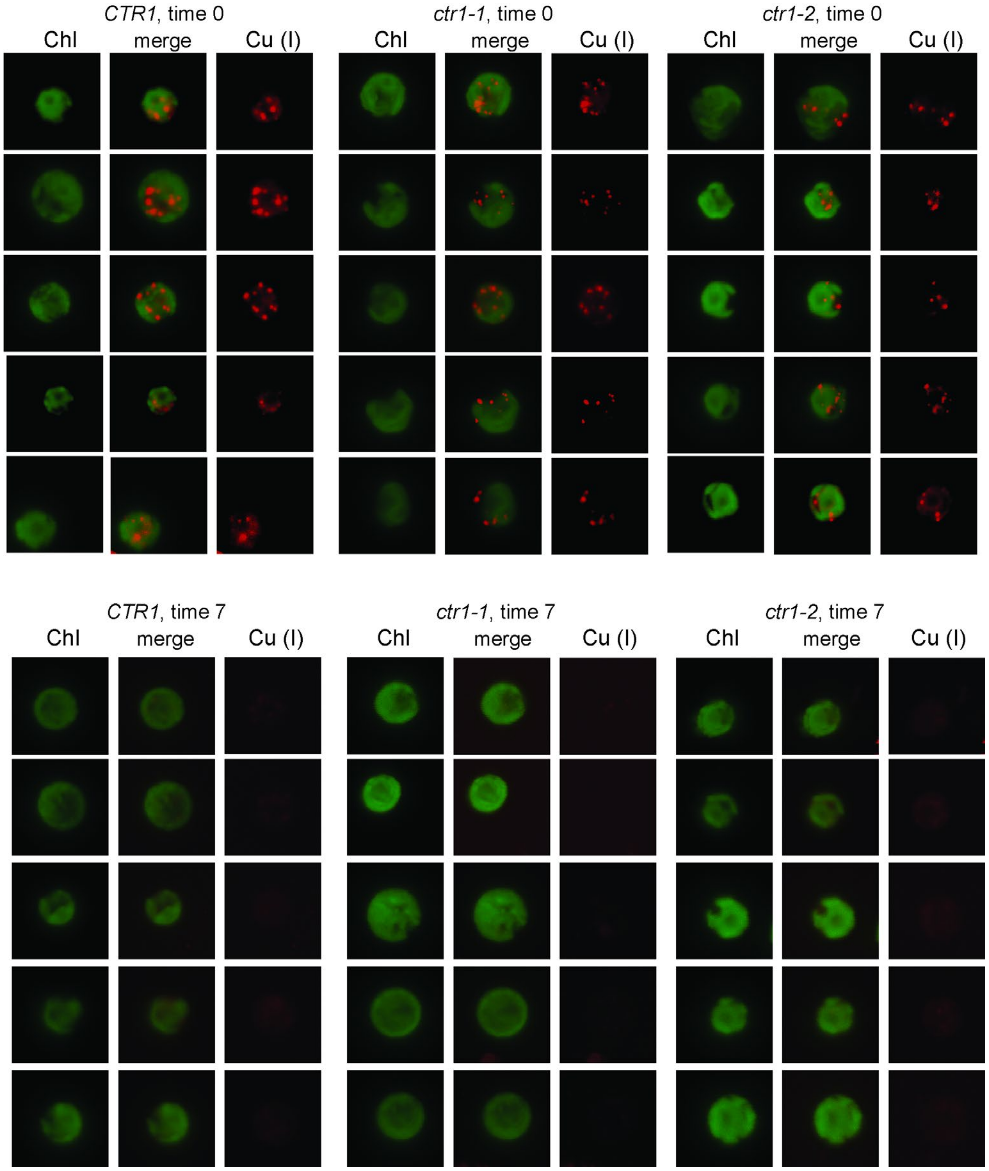


**Supplemental Figure 12. Copper accumulated in zinc deficient *ctr1* mutants**. Figure shows all *ctr1* cells that were imaged in experiments shown and described in Figure 9.

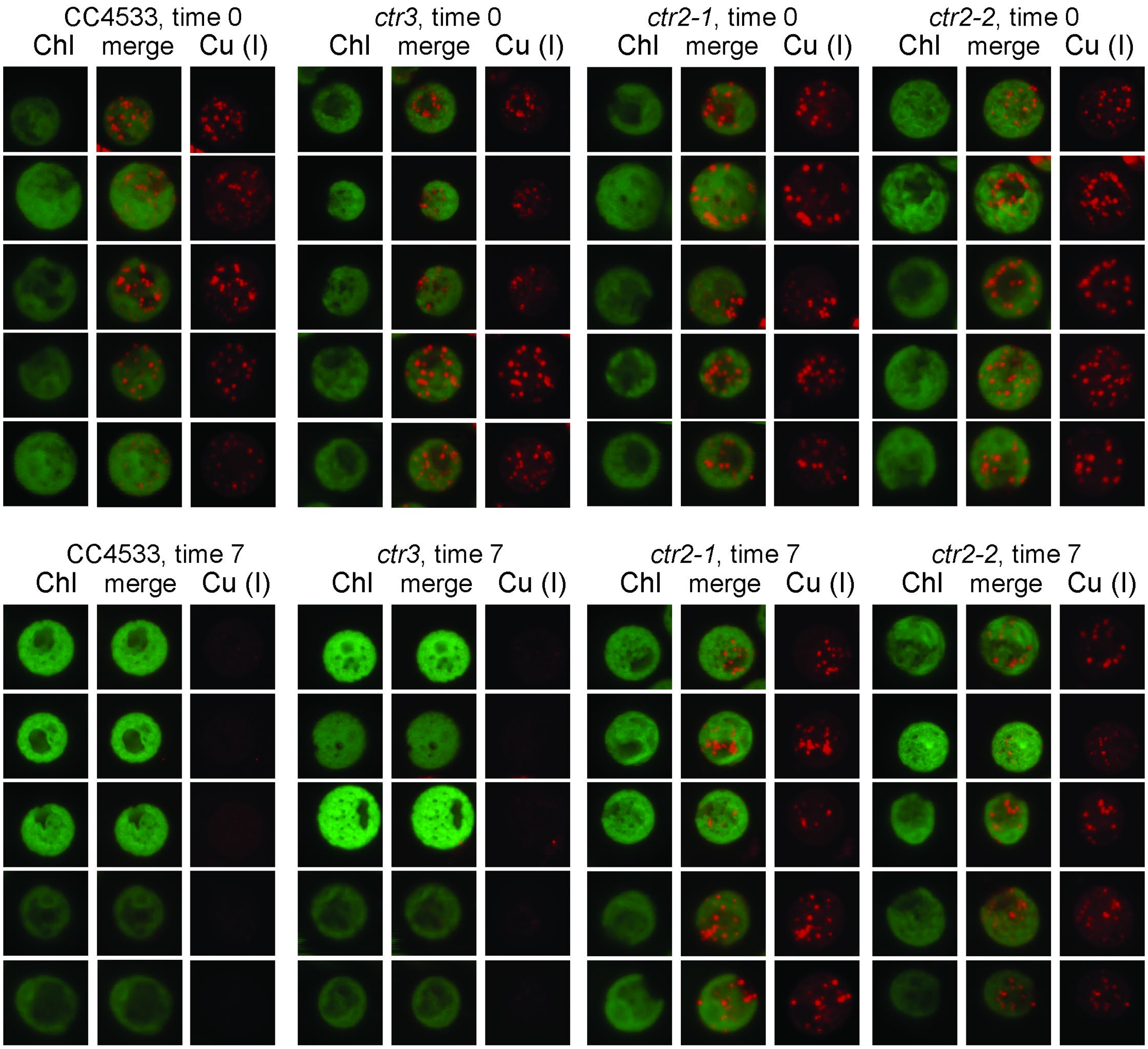


**Supplemental Figure 13. Copper accumulated in zinc deficient *ctr2* and *ctr3* mutants**. Figure shows all *ctr2* and *ctr3* cells that were imaged in experiments shown and described in Figure 9.
